# Symbiotic signalling is at the core of an endophytic *Fusarium solani*-legume association

**DOI:** 10.1101/740043

**Authors:** Skiada Vasiliki, Marianna Avramidou, Paola Bonfante, Andrea Genre, Kalliope K. Papadopoulou

**Affiliations:** Department of Biochemistry and Biotechnology, University of Thessaly, Biopolis, Larissa, 41500, Greece; Department of Life Sciences and Systems Biology, University of Torino, Torino, 10125, Italy

## Abstract

Legumes interact with a wide range of microbes in their root system, ranging from beneficial symbionts to pathogens. Symbiotic rhizobia and arbuscular mycorrhizal glomeromycetes trigger a so-called common symbiotic signalling pathway (CSSP), including the induction of nuclear calcium spiking in the root epidermis. In our study, the recognition of an endophytic *Fusarium solani* strain K in *Lotus japonicus* induced the expression of *LysM* receptors for chitin-based molecules, CSSP members and CSSP-dependent genes in *L. japonicus*. In *LysM* and CSSP mutant/RNAi lines, root penetration and fungal intraradical progression was either stimulated or limited while FsK exudates are perceived in a CSSP-dependent manner, triggering nuclear calcium spiking in epidermal cells of *Medicago truncatula* Root Organ Cultures. Our results corroborate that the CSSP is a more common pathway than previously envisaged, involved in the perception of signals from other microbes beyond the restricted group of symbiotic interactions *sensu stricto*.

## Introduction

Plants encounter diverse microorganisms at the root-soil interface, some of which invade the root system and establish detrimental or beneficial associations (Zipfel and Oldroyd, 2017). In this scenario, legumes are unique in their ability to establish two types of mutualistic symbioses: the arbuscular mycorrhizal (AM) and rhizobium-legume (RL) symbiosis. For both interactions, a few plant-directed microbial signals have been characterized so far, and all of them are water soluble chitin-based molecules. Rhizobia produce Nod Factors (NFs), lipo-chitooligosaccharides (LCOs) that are recognized by specific receptor-like kinases (RLKs) (Amor et al., 2003) (Limpens et al., 2003) (Madsen et al., 2003) (Murakami et al., 2018) (Radutoiu et al., 2003), whereas AM fungi release both Nod factor-like Myc-LCOs (Maillet et al., 2011) and short-chain chitin oligomers (Myc-COs) (Genre et al., 2013). The perception of chitin-related molecules in plants is mediated by Lysin-motif (LysM)-RLKs (Antolín-Llovera et al., 2014). This is also the case for Nod factors and Myc-LCOs (Fliegmann et al., 2013) (Maillet et al., 2011), whereas specific receptors for Myc-COs remain to be characterized.

Downstream of receptor activation, the intracellular accommodation of both types of symbionts is mediated by a so-called common symbiosis signalling pathway (CSSP) that in *Lotus japonicus* includes eight genes: the plasma membrane-bound RLK *Lj*SYMRK (*M*tDMI2 in *Medicago truncatula*), three nucleoporins (NUP85, NUP133, NENA), the nuclear envelope located cation channels *Lj*CASTOR, and *Lj*POLLUX (*Mt*DMI1, orthologous to POLLUX), the nuclear calcium and calmodulin-dependent protein kinase *Lj*CCaMK (*Mt*DMI3), the CCaMK phosphorylation substrate *Lj*CYCLOPS (Kistner et al., 2005) (Charpentier et al., 2008) (Genre and Russo, 2016). Downstream the CSSP lie nodulation/mycorrhization specific transcription factors such as *Lj*NSP1 and *Lj*NSP2 (Catoira et al., 2000) (Delaux et al., 2013) (Kalo et al., 2005) (Maillet et al., 2011), and early nodulin genes, like *LjENOD2* and *LjENOD40* (Glyan’ko, 2018) (Ferguson and Mathesius, 2014) (Takeda et al., 2005). A central element of the CSSP is the triggering of repeated oscillations in nuclear calcium concentration (spiking). In fact, mutants in all CSSP members acting upstream of CCaMK are defective for calcium spiking (Harris et al., 2003) (Miwa et al., 2006) (Parniske, 2008), and the nuclear-localized CCaMK is considered to act as a decoder of the calcium signal, which is translated into phosphorylation events (Yano et al., 2008). Consequently, the induction of nuclear calcium spiking has become of common use as a reliable reporter of CSSP activation by symbiotic microbes or their isolated signals (Wais et al., 2000) (Sieberer et al., 2009) (Chabaud et al., 2011) (Genre et al., 2013) (Charpentier and Oldroyd, 2013). In particular, this approach has been used for the study of AM fungal signalling in both legume and non-legume hosts, using either whole plants or root organ cultures (ROCs) expressing nuclear-localized calcium-sensing reporters, such as cameleon proteins (Chabaud et al., 2011) (Genre et al., 2013) (Sun et al., 2015) (Carotenuto et al., 2017).

A role for the CSSP in non-symbiotic interactions, has been investigated by many research groups: some propose an (at least partial) involvement (Sanchez et al., 2004) (Sanchez et al., 2005) (Weerasinghe et al., 2005) (Fernandez-Aparicio et al., 2010) (Wang et al., 2012) (Zgadzaj et al., 2015) (Zgadzaj et al., 2019), while others do not (Banhara et al., 2015) (Huisman et al., 2015). Nevertheless, little is known about the triggering of nuclear calcium oscillations in such associations, whereas compromised colonization phenotypes by a non-symbiont has not been reported, thus far, for a CSSP mutant.

FsK is a beneficial endophytic isolate of tomato, protecting the plant against root and foliar pathogens (Kavroulakis et al., 2007), spider mites (Pappas et al., 2018), zoophytophagous insects (Garantonakis et al., 2018), and drought (Kavroulakis et al., 2018). We have recently shown that FsK efficiently colonizes legume roots and demonstrates great plasticity throughout colonization (Skiada et al., 2019). Commonalities in the host cell responses between legumes-FsK, and to both symbiotic and pathogenic interactions (Genre et al., 2009) (Skiada et al., 2019), prompted us to investigate the role of the CSSP in this endophytic legume association.

We herein used gene expression analysis and phenotypic screening of mutant/RNAi lines to investigate the role of certain *LysM* receptors in the *Lotus*-FsK interaction. We furthermore demonstrate that the expression of *L. japonicus* markers of AM and/or RL symbiosis is partially altered in the presence of FsK, while two CSSP mutants, *ccamk* and *cyclops*, display a delay in fungal colonization. Lastly, we show that the endophyte exudates triggered CSSP-dependent nuclear calcium spiking in the outer root tissues of *M. truncatula* ROCs, which is also observed for exudates from other root-interacting fungi. Overall, our results show that multiple *LysM* receptors participate in FsK recognition and suggest that the core of the CSSP is also involved in the perception of diffusible signals from non-mycorrhizal fungi.

## Results

### FsK inoculation affects the expression of CSSP members and CSSP-regulated genes

Early FsK colonization of legume roots triggers a number of host cell responses (cytoplasm/ER accumulation, nuclear movement and membrane trafficking at contact site) that are also known to occur in both symbiotic and pathogenic interactions (Skiada et al., 2019). This prompted us to investigate potential similarities with symbiosis-related plant gene expression. We, therefore, analyzed the expression of key CSSP components, such as *LjCASTOR*, *LjCCaMK* and *LjCYCLOPS*, as well as CSSP-regulated symbiosis markers, namely *LjNSP1*, *LjNSP2*, *LjIPN2*, *LjENOD40 and LjENOD2*. Their regulation was investigated at specific time points during *Lotus*-FsK interaction, alongside fungal colonization levels (Supplementary Figure 1, 10). The expression of *LjCCaMK* was marginally but significantly upregulated upon FsK inoculation at very early stages of the interaction (1 dpi), reaching a 2.2-fold upregulation compared to controls at 12 dpi. By contrast, no significant difference was recorded in the expression of *LjCASTOR* and *LjCYCLOPS*, or the transcription factors *LjNSP1* and *LjNSP2*, or *LjIPN2* (Interacting Protein of NSP2) acting downstream the CSSP, in control and inoculated plants at any time point (Supplementary Figure 1, 10).

We then tested whether the expression of the symbiosis marker gene *LjENOD40* was altered in our *Lotus*-FsK interaction system. We focused on *LjENOD40-1* gene, which is strongly upregulated at very early stages of rhizobium infection, in contrast to the mature nodule marker *LjENOD40-2* (Kumagai et al., 2006). A statistically significant induction of *LjENOD40-1* was observed at 2, 4 and 6 dpi (1.6-, 1.9-, 1.7-fold, respectively) in inoculated compared to non-inoculated plants. *ENOD40* expression coincides with maximum levels of intraradical fungal accommodation (recorded at 4 dpi) (Supplementary Figure 10). By contrast, we recorded no differences between control and inoculated plants in the expression levels of our second nodulation marker gene, *ENOD2*, which is expressed in legumes during rhizobial and AM infection and nodule morphogenesis, but not upon fungal pathogen infection (Franssen et al., 1987) (van de Wiel et al., 1990) (van Rhijn et al., 1997).

To summarize, our analyses revealed that FsK colonization was associated with the upregulation of a subset of known symbiosis markers over the course of 12 days post inoculation.

### FsK colonization is reduced in CSSP mutants compared to wt plants

Based on our gene expression results, we investigated the phenotypic differences in FsK colonization of *L. japonicus* CSSP mutants compared to wt plants (Figure 1). We selected mutant lines impaired in CSSP genes acting upstream (*symRK-1*, *castor-1*) and downstream of nuclear calcium spiking (*ccamk-1*, *cyclops-1*). FsK intraradical colonization was quantified via qPCR at 4 and 8 dpi (Figure 1Β), when roots are well-colonized by the fungus under our experimental conditions (Supplementary Figure 10). The colonization of *symRK-1* and *castor-1* mutants was similar to that of wt plants. By contrast, a significant reduction in fungal colonization was observed in both *ccamk-1* (75.17%) and *cyclops-1* (68.82%) compared to wt plants at 4 dpi. A comparable reduction in fungal intraradical colonization persisted at 8 dpi only in *cyclops-1* plants (45.77%).

**Figure 1.**
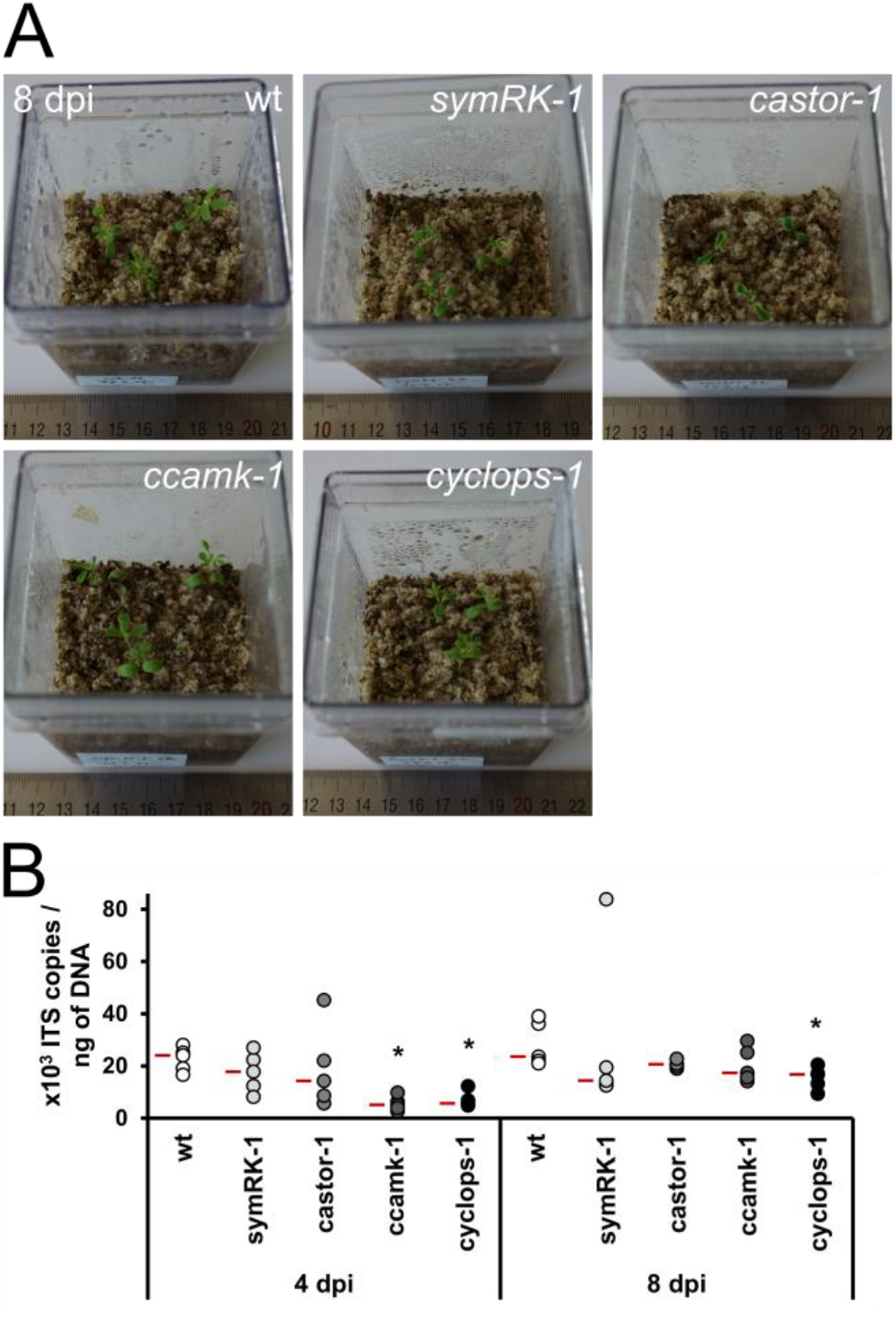
Quantification of FsK colonization in *L. japonicus* CSSP mutants. **A** FsK inoculated *Lotus japonicus* wt and CSSP mutants at the last time point of the experiment (8 dpi). **B** Absolute quantification of FsK ingress within *L. japonicus* root tissues in wt and CSSP mutants (*symRK-1*, *castor-1*, *ccamK-1*, *cyclops-1*), via qPCR, using primers specific for a fragment of *Fusarium ITS* gene. Tissues were harvested at 4 and 8 dpi. *ITS* gene copy numbers are normalized to ng of total DNA extracted from the root tissue. Data are presented as a dotplot. Each dot represents a single biological replicate. 5 biological replicates were assessed for each genotype. Each replicate consists of 3 individual plants. Median values are presented in red. The experiment was repeated twice with similar results. Asterisks represent statistically significant differences between wt and the respective mutant line at the 0.05 level (Student’s t-test).

In conclusion, our phenotypic analysis revealed a significant reduction in FsK colonization only for CSSP mutants for genes that act downstream of nuclear calcium spiking, and the reduction was more evident at earlier time points of the interaction.

### Fungal exudates trigger nuclear calcium spiking in wt *M. truncatula* ROC epidermis

Based on the colonization phenotype in CSSP mutant lines, we then investigated nuclear calcium signals in response to FsK exudate treatment. To this aim, we used wt *Medicago truncatula* ROCs expressing the nuclear localized cameleon reporters (NupYC2.1), thus allowing FRET based imaging analysis of nuclear calcium responses (Chabaud et al., 2011) (Carotenuto et al., 2019).

We focused on epidermal cells in an area 1-2 cm above the root tip harboring few trichoblasts, because this root zone in model legumes is susceptible to colonization by FsK (Skiada et al., 2019) and is also reported to show the strongest calcium spiking responses to AM fungal exudate (Chabaud et al., 2011). As additional controls, we extended our analysis to include other compatible legume colonizers: the pathogenic *Fusarium oxysporum* f.sp. *medicaginis* (Snyder and Hansen, 1940), and *Piriformospora (Serendipita) indica* (Hayes et al., 2014) (Ramírez-Suero et al., 2010), an endophytic fungus with a versatile lifestyle (Verma et al., 1998) (Deshmukh et al., 2006) (Lahrmann et al., 2013).

Exudates from all tested fungi triggered nuclear calcium oscillations in wt *M. truncatula* epidermal cells (Figure 2, Supplementary Figure 2; details about the biological replicates and nuclei screened are presented in Supplementary Table 1). A higher number of cells responded to FsK (70.11%) or Fom (80.95%), compared to *P. indica* exudate (43.59%; Figure 3A). In addition, although the median number of spikes generated in active cells over 30 minutes of recording was comparable for all exudates (5 peaks for FsK and Fom; 4 peaks for *P. indica*; *p*-value 0.245) (Figure 3B), when we analysed the distribution of peak numbers among the fungal exudates tested in all cells, we recorded differences among peak numbers generated by *P. indica* and FsK or Fom, respectively (p-value <0.005 for each pairwise comparison). Thus, differences in the pattern of calcium spikes, expressed as the number of spikes per cell over a period were present.

**Figure 2.**
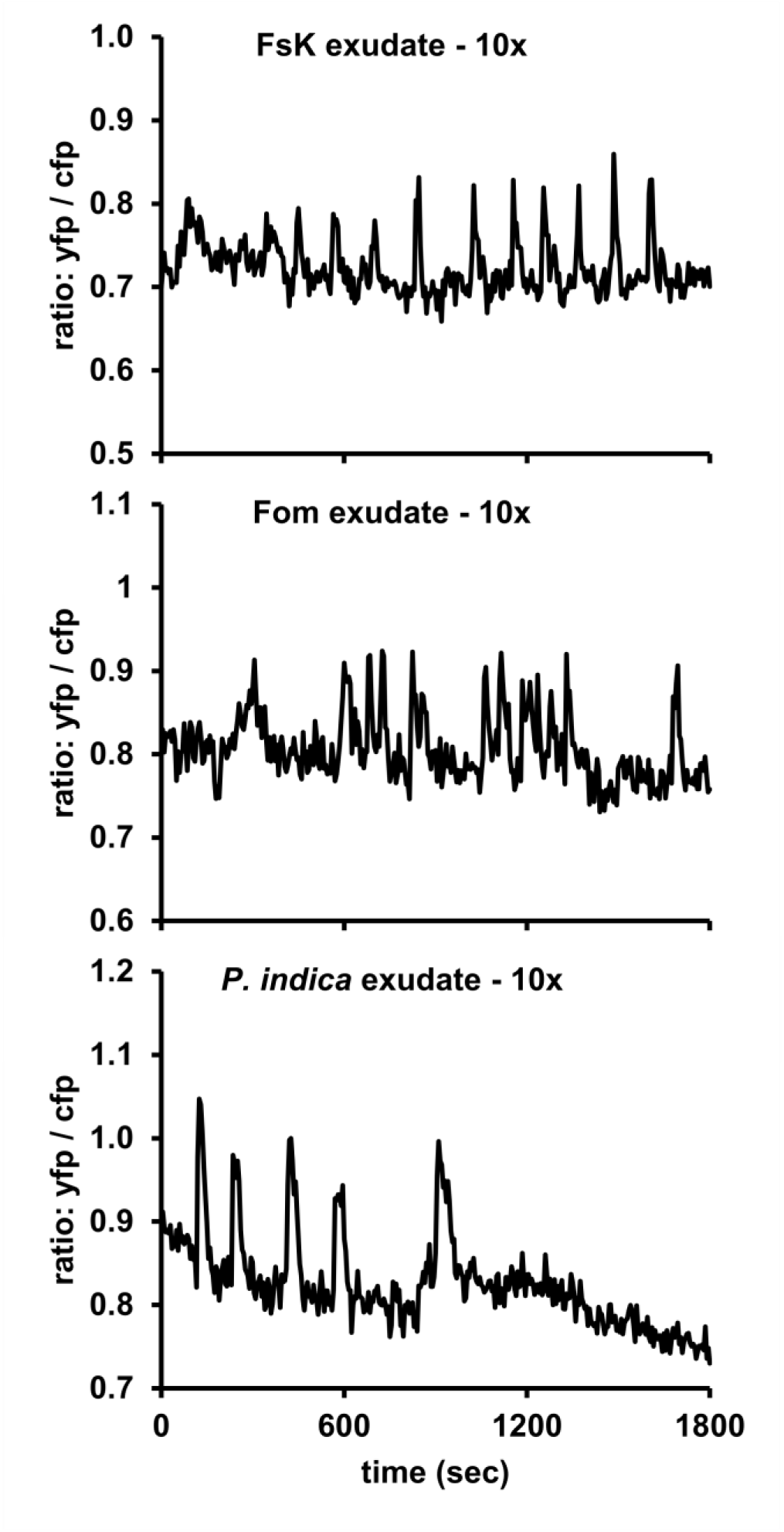
Exudates produced by fungi induce changes in nuclear calcium levels imaged using cameleon *Mt* ROCs. Calcium spiking responses were recorded in a low-trichoblast-region located 10-20 mm from the root tip. *M. truncatula* wt ROCs expressing the NupYC2.1 cameleon were used for the bioassay. Fungal exudates derived from FsK, Fom, or *P. indica* were used as elicitors. Representative plots showing changes in nuclear calcium levels after treatment with 10x concentrated FsK, Fom, or *P. indica* exudate, in a 30 min (1800 sec) recording. Graphs show ratios of YFP:CFP fluorescence over time. The respective changes in fluorescence over time are presented in graphs of Supplementary Figure 2.

**Figure 3.**
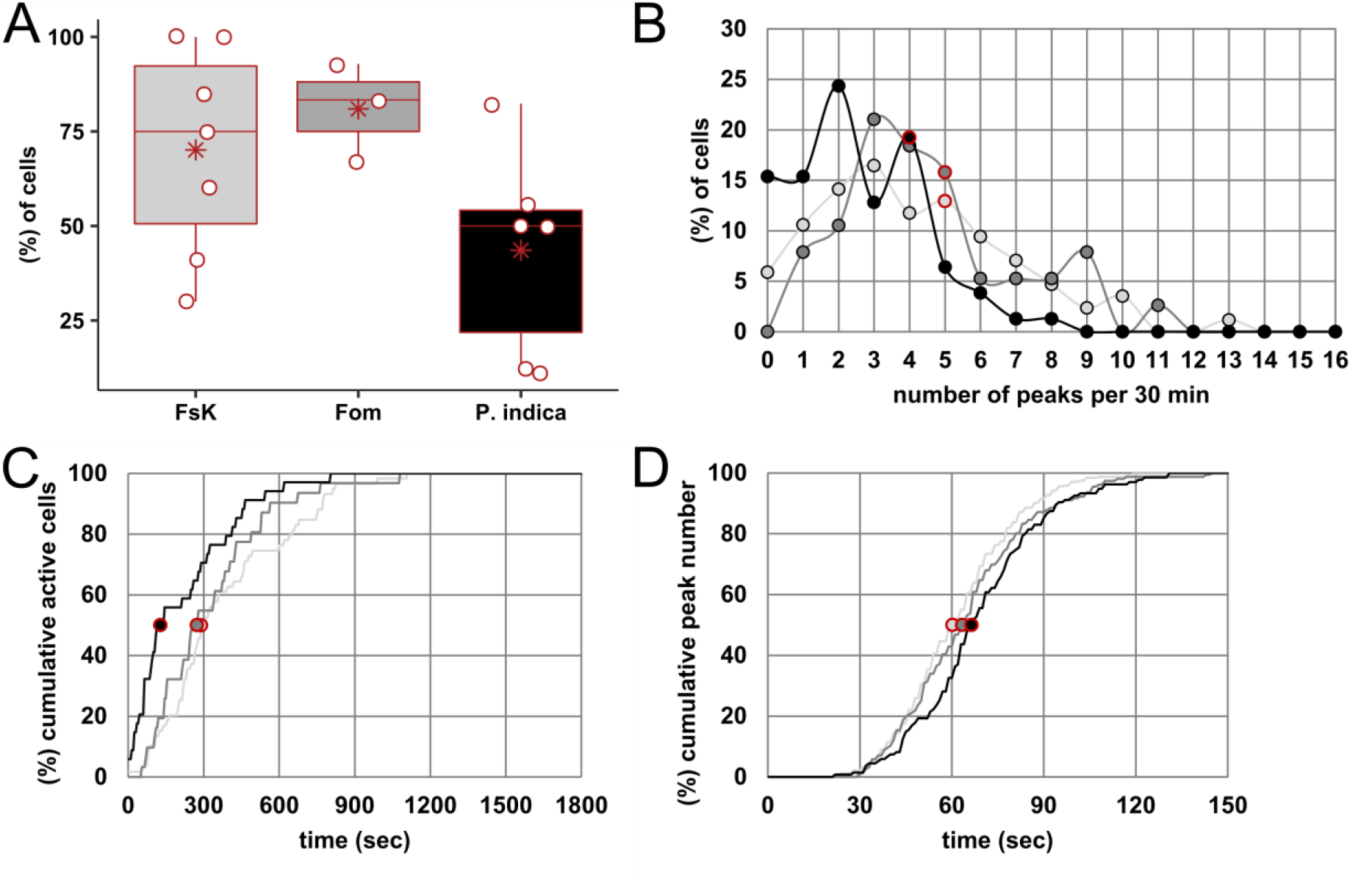
Analysis of fungal exudate-triggered nuclear calcium responses in *Mt* wt ROCs epidermis. Calcium spiking responses were recorded in a low-trichoblast-region located 10-20 mm from the root tip. *M. truncatula* wt (ROCs) expressing the NupYC2.1 cameleon were used for the bioassay. Fungal exudates derived from FsK, Fom, or *P. indica* were used as elicitors. **A** Percentage of responsive cells (peak number>2), in 30 min recordings post fungal exudate application. Data are presented as a dotplot in combination with a boxplot for each exudate. Each dot represents a single biological replicate (individual lateral root segments derived from ROCs). Median values are presented as red line within the boxplot, whereas average values are presented as a red asterisk. No statistically significant difference was recorded in average percentages of cells showing calcium spikes in response to FsK/FoM/P. indica exudate elicitation (one-way anova). **B** Histogram exhibiting the distribution of fungal exudate-induced calcium spiking responses in all nuclei examined (active and non-active cells). Dots represent the frequency of peak number (presented in continuous intervals of 1 peak) in a 30 min recording period. Median peak number values calculated only in active cells are also presented in red. Two-sample Kolmogorov Smirnov (K-S) test was performed to compare the distribution of peak numbers generated in response to the three exudates. Distributions were compared in a pairwise manner, under the null hypothesis that populations are drawn from the same distribution. When all cells (active and non-active) were included in the analysis: FsK-*P. indica* and Fom-*P. indica* comparison showed that data are drawn from different distributions (p<0.005 for both comparisons). **C** Cumulative probability line plots showing, for each exudate (depicted by different line colour), the percentage of active cells as functions of the delay time (in sec) recorded prior to spiking initiation. Lag phases were measured manually for all active nuclei examined per elicitor. Lag phase represents the time intervening between the recording initiation and the first recorded spike for each responsive nucleus. Median delay time values are also presented in red. Two-sample K-S test was then performed to compare the distribution of lag phases recorded in response to each elicitor. Distributions were compared in a pairwise manner, under the null hypothesis that populations are drawn from the same distribution. Pairwise comparisons among FsK-*P. indica* and Fom-*P. indica* showed that data are drawn from different distributions (p<0.005 and p<0.05, respectively). **D** Cumulative probability line plots showing, for each elicitor (depicted by different line colour), the percentage of active cells as functions of the spike width (in sec). Spike widths were measured manually for all peaks recorded and for each fungal exudate applied. Median spike width values are also presented in red. Two-sample K-S test was then performed to compare the distribution of spike width values recorded in response to each elicitor. Distributions were compared in a pairwise manner, under the null hypothesis that populations are drawn from the same distribution. Pairwise comparisons among FsK-*P. indica* showed that data are drawn from different distributions (p<0.005). Light grey/dark grey/black lines in panels B, C, D: frequencies for FsK/FoM/*P. indica* exudate-elicited calcium spiking responses, respectively. The number of biological replicates/nuclei/peaks assessed for each elicitor and for each analysis is indicated in Supplementary Table 1.

It has been postulated that information in calcium-mediated signals can be encoded in the amplitude and frequency of the spiking, duration of the response as well as tissue specificity, and this information is responsible for the induction of specific responses (McAinsh and Hetherington, 1998) (Vadassery et al., 2009). Thus, we analyzed (Figure 3C) the oscillatory responses generated by our fungal exudates (Figure 2) by manually calculating the interval between the treatment and the occurrence of the first spike in active cells (lag phase). Differences in median lag phase values of active cells examined per exudate were sharp (127.96 sec for *P. indica*, 290.32 sec for FsK and 273.12 sec for Fom), and the distribution of lag phase values were statistically different for induced calcium spiking between *P. indica*-FsK and *P. indica*-Fom (p-value <0.005 and p-value <0.05, respectively).

We also measured the width of each spike in all active cells, estimated as the difference in seconds between the first time point of the upward phase (elevation from the baseline - steady state) and the last time point of the downward phase (return to the baseline – steady state) for each spike. Spikes generated by the 3 fungal exudates had comparable median values (60.22 sec for FsK, 63.38 sec for Fom, 66.47 sec for *P. indica;* as calculated in the total number of active cells). Analysis of the distribution of spike width values recorded in response to the three exudates, revealed differences among FsK and *P. indica* (p<u>-value</u> <0.005).

Lastly, we analysed the waiting time, i.e. the interval between subsequent peaks. The histogram in Supplementary Figure 4 illustrates the distribution of waiting time autocorrelation values in our populations of spiking profiles. A high number of cells in all treatments demonstrated negative autocorrelation values, indicating that spiking responses to all fungal exudates are in most cases irregular, in analogy to what was described for AM fungal-induced spiking profiles and in contrast with the mostly regular rhizobium-induced calcium spiking signals in the same experimental system (Russo et al., 2013). Statistical analysis showed no dependency between the distribution of waiting time autocorrelation values and the fungal exudate tested.

**Figure 4.**
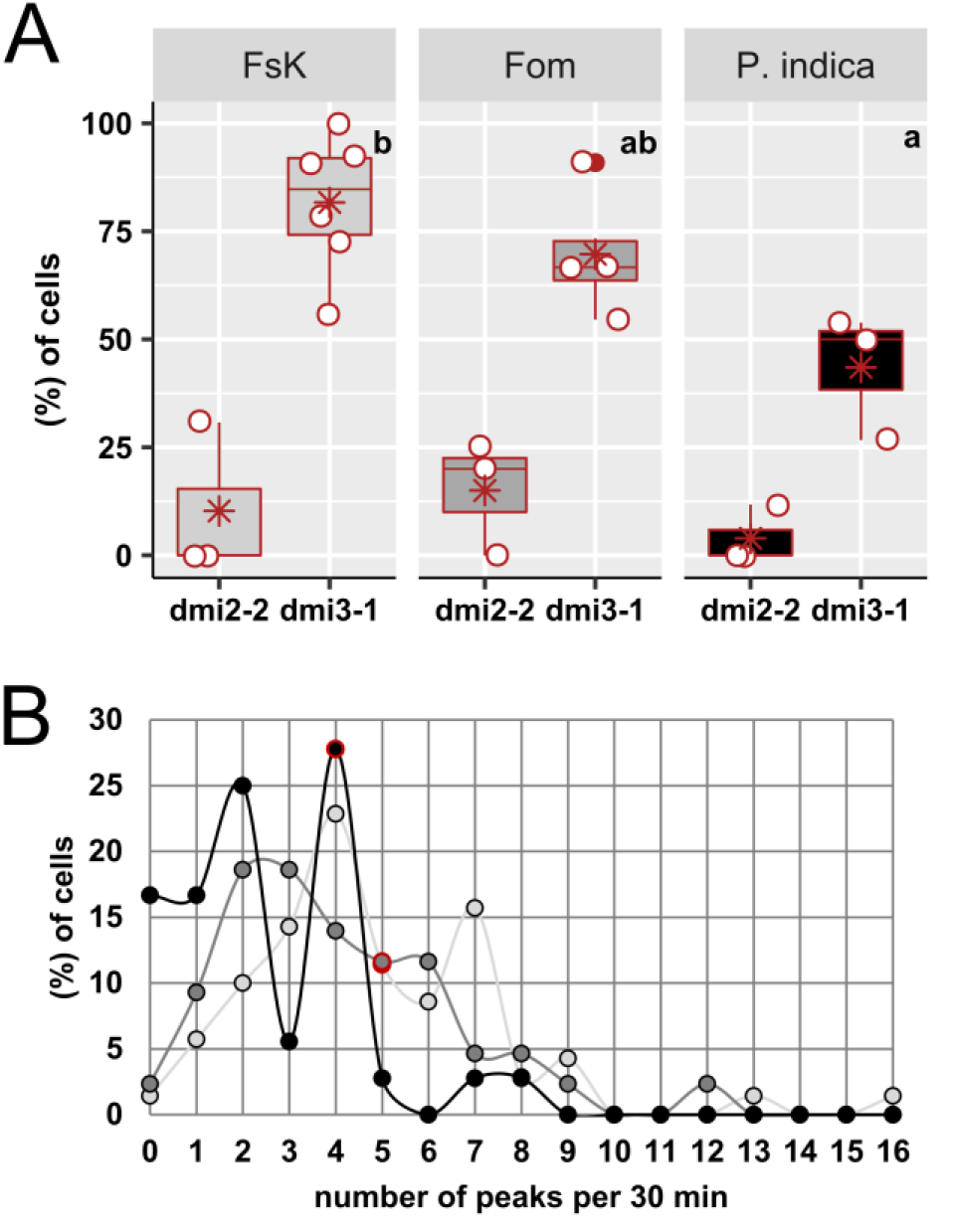
Analysis of fungal exudate-triggered nuclear calcium responses in *Mt dmi2-2* and *dmi3-1* ROCs epidermis. Calcium spiking responses in a low-trichoblast-region located 10-20 mm from the root tip. *M. truncatula dmi2-2* and *dmi3-1* mutant lines (ROCs) expressing the NupYC2.1 cameleon were used for the bioassay. Fungal exudates derived from FsK, Fom, or *P. indica* were used as elicitors. **A** Percentage of responsive cells (peak number>2) in the *dmi2-2* and *dmi3-1* mutant background, in 30 min recordings. Data are presented as a dotplot in combination with a boxplot for each mutant line/exudate. Each dot represents a single biological replicate (individual lateral root segments derived from ROCs). Median values are presented as red line within the boxplot, whereas average values are presented as a red asterisk. Note that an outlier is presented in dmi3-1 lines for Fom exudate (presented as a red dot). Outlier value was not removed from subsequent analysis as it was considered a result of biological variation. Different letters indicate statistically significant differences in percentages of *dmi3-1* responding cells to each elicitor (one-way Anova performed in biological replicates, followed by Tukey’s post-hoc test). **B** Histogram exhibiting the distribution of fungal exudate-induced calcium spiking responses in nuclei of *dmi3-1* ROC segments (active and non-active cells). Dots represent the frequency of peak number (presented in continuous intervals of 1 peak) in a 30 min recording period. Median peak number values calculated only in only active cells are presented in red. Two-sample (K-S) test was then performed to compare the distribution of peak numbers generated in response to fungal exudates. Distributions were compared in a pairwise manner, under the null hypothesis that populations are drawn from the same distribution. When all cells (active and non-active) were included in the analysis, FsK-*P. indica* comparison showed that data are drawn from different distributions (p<0.001). Light grey/dark grey/black lines: frequencies of peak number values of FsK/FoM/*P. indica* exudate-elicited calcium spiking responses in panel B, respectively. The number of biological replicates/nuclei assessed for each elicitor/ROC line is indicated in Supplementary Table 1.

Altogether our analysis revealed that the conditions (i.e. constituents of the exudates) that determined the distribution of 1) peak number generated over a certain period of time, 2) lag phase prior to spiking initiation, and, 3) peak width of individual traces in response to the 3 elicitors examined, were not the same. More specifically, *P. indica-*induced calcium spiking had the most profound differences in comparison to *Fusaria*-induced calcium spiking. Pairwise comparison among the two *Fusaria* examined, revealed no differences in any of the parameters examined herein.

### Nuclear calcium spiking is abolished in *Mtdmi2-2* but not in *Mtdmi3-1* epidermis

We then extended our investigation of nuclear calcium signals in ROC epidermal cells of mutants for either *MtDMI2 (dmi2-2*) and *MtDMI3* (*dmi3-1*), homologs to *LjSYMRK* and *LjCCAMK*, respectively (Figure 4, Supplementary Figure 5). In *dmi2-2*, very few cells were active in any treatment (10.26% for FsK, 15% for Fom, 3.92 % for *P. indica* exudate). By contrast, nuclear calcium spiking was retained in *dmi3-1* upon stimulation by the fungal exudates tested (Figure 4A). Furthermore, FsK exudate induced nuclear spiking in a higher percentage of *dmi3-1* cells (81.68%) compared to wt (70.11%). The opposite was observed for Fom exudate (69.70% for *dmi3-1* vs 80.95% for wt), whereas approximately the same percentage of responding cells was recorded in *dmi3-1* and wt treated with *P. indica* exudate (43.50% for *dmi3-1* vs 43.59% for wt) (Figure 4A).

**Figure 5.**
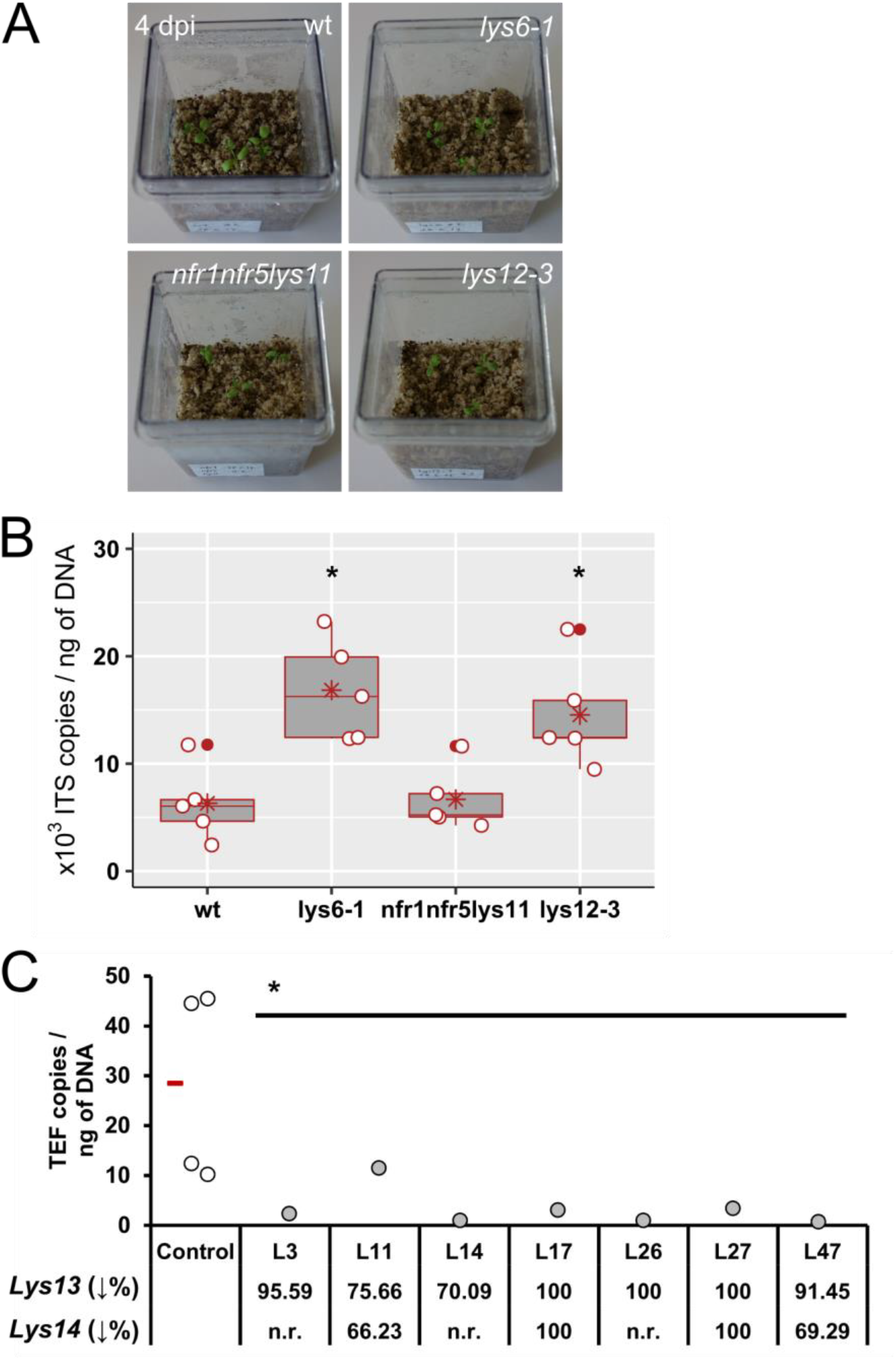
Quantification of FsK abundance in roots of *L. japonicus* mutants impaired in *LysM* receptors, and in *LysM13*/*14* RNAi lines. **A** FsK inoculated *L. japonicus* wt and *LysM* mutants at the last time point of the experiment (4 dpi). **B** Absolute quantification of FsK ingress within *L. japonicus* root tissues in wt and *LysM* mutants (*lys6-1, nfr1nfr5lys11, lys12-3*), via qPCR, using primers specific for a fragment of *Fusarium ITS* gene. Tissues were harvested at 4 dpi. *ITS* gene copy numbers are normalized to ng of total DNA extracted from wt and mutant root tissues, respectively. Data are presented as a dotplot. Each dot represents a single biological replicate. Median values are presented in red. 5 biological replicates were assessed for each genotype. Each replicate consists of 3 individual plants. Asterisks represent statistically significant differences between wt and the respective mutant lines at the 0.05 level (Student’s t-test). Note that there are outlier values in wt, *nfr1nfr5lys11*, and *lys12-3* lines (presented as a red dot). Outlier values were not removed from subsequent analysis as they were considered a result of biological variation. **C** Absolute quantification of FsK abundance within control lines (presented as average values of 4 independent plants) transformed with an empty T-DNA vector, and within 7 independent hairy root lines transformed with a *Lys13*/*14* RNAi silencing construct (L3-L47). Quantification was performed via qPCR, using primers specific for a fragment of *Fusarium* TEF gene. Tissues were harvested at 5 dpi. TEF gene copy numbers are normalized to ng of total DNA extracted from each root tissue sample. Data are presented as a dotplot. For the control, each dot represents a single biological replicate. Median value is presented in red. Each replicate consists of an individual plant. 4 independent plants were assessed. For the *LysM13*/*14* RNAi lines, each dot represents the mean of 2 technical replicates. Mean FsK abundance value (average TEF gene copy numbers) of all *Lys13*/*14* RNAi lines is significantly lower than that of controls at the 0.05 level (Student’s t-test). The % reduction in *Lys13* or *Lys14* relative transcript levels in RNAi lines in comparison to relative transcript levels in controls (as quantified via gene expression analysis; presented in Supplementary Figure 9) is presented at the lower part of the graph. n.r., no reduction in respective relative transcript levels.

We then analysed the distribution of peak number in *dmi3-1* cells generated by the three fungal exudates (Figure 4B). The median number of spikes for active cells was within similar ranges in all exudates (similar median values to those obtained in the wt lines) (Figure 4B). On the other hand, a significant association was recorded in the peak distribution in total cells (active and non-active) among FsK and *P. indica* exudate (p-value <0.001), indicating again differences in the number of spikes per cell over a period as observed in wt.

In conclusion, all fungal exudates examined generated intense and repeated nuclear calcium oscillations (spiking). Their activation was dependent on *DMI2* (a member of the CSSP acting upstream of calcium oscillations) but not on *DMI3* (acting downstream of calcium signalling), suggesting that the core of the CSSP that transduces AM fungal, rhizobial and actinorrhizal signals (Barker et al., 2017) is also involved in the perception of signals from all three non-mycorrhizal fungi.

### The biological activity of *Fusarium* exudates is affected by chitinase and heat treatment

In an attempt to determine the nature of the factors that triggered nuclear calcium spiking by FsK, we hypothesized that if the triggering compounds were chitin-based molecules, their enzymatic cleavage with chitinase would lead to a decrease in the spiking response (Genre et al., 2013) (Chabaud et al., 2016). For this quest, we also included in the analysis the exudates of Fom, since it belongs in the *Fusarium* genus, and the two fungal strains produced similar calcium spiking responses in wt and *dmi3-1* cells. Supplementary Figure 6 shows that the percentage of epidermal cells responding to chitinase-treated FsK exudate dropped significantly by 62.86%, and an analogous 44.68% reduction was observed upon chitinase treatment of Fom exudate, compared to their respective control experiments with untreated exudates. As a control for the enzymatic activity of the chitinase solution, we used short-chain COs that are known to elicit calcium spiking in atrichoblasts (Genre et al., 2013). Pre-treatment of 10^−6^ M CO5 with chitinase resulted in partial abolishment of nuclear calcium oscillatory response in our experiments (32.12% reduction; Supplementary Figure 7).

As a second step, we examined the stability of the two exudates under heat treatment. To this aim, we autoclaved FsK and Fom exudates at 120°C for 20 min. Again, the reduction in the percent of cells responding to heat-treated exudates as compared to the non-treated ones was more severely affected for FsK (55.92 %) than Fom (14.22 %). A prolonged heat treatment of 60 min did not reduce further the activity of FsK exudates (56.70 % reduction in responding epidermal cells).

In short, our tests suggested that the active molecules present in FsK and Fom exudates include both chitinase- and heat-sensitive compounds. Furthermore, the observed differences in the reduction of the activity between heat-treated FsK and Fom exudates suggests that the molecules that trigger a comparable nuclear calcium spiking pattern are different in the two fungal species.

### Reaching the pathway backwards: involvement of LysM-RLKs in *Lotus japonicus* response

Our results on FRET-based analysis of nuclear calcium changes in wt ROC cells upon chitinase-treated FsK exudate, prompted us to investigate the role of members of the *LysM* RLK gene family in *Lotus*-FsK interaction. We focused on genes that are regulated upon chitin/fungal treatment (Fuechtbauer et al., 2018) (Lohmann et al., 2010) (Rasmussen et al., 2016), and followed their transcriptional regulation over time during *Lotus* root colonization by FsK. More specifically, we investigated the transcript levels of 6 *LjLys* genes: *Lys6*, *Lys11*, *Lys12*, *Lys13*, *Lys14*, *Lys20* (Supplementary Figure 8, 10). Four *LysM* genes were transcriptionally regulated during the interaction with FsK: *Lys6* had a statistically significant upregulation (1.5-fold) at 4 dpi, which correlates with maximal fungal presence within plant tissues at this time point (Supplementary Figure 10); *Lys13* was upregulated from 2 dpi onwards, and *Lys14* is upregulated only at late stages of the interaction (12 dpi). *Lys12* and *Lys20* expression was not altered in the presence of FsK, while *Lys11* transcript levels were very low in both control and inoculated tissues, under our experimental conditions.

To further investigate the possible role of differentially expressed *LysM* genes in *Lotus*-FsK interaction, we compared FsK colonization levels between wt and mutant lines impaired in *LysM* genes that were also, transcriptionally affected. *Ljlys6-1* displayed significantly higher FsK colonization levels at 4dpi compared to wt plants. Since mutant lines for *LjLys13* and *LjLys14* were not available, we opted for the generation of silenced hairy-root lines, to examine the role of these genes in FsK colonization. We introduced an RNA-hairpin construct targeting both *Lys13* and *Lys14* into *L. japonicus* roots, using the *Agrobacterium rhizogenes*-mediated hairy root transformation method. We obtained 7 individual plants showing 70% to 100% reduction in *Lys13* relative transcript levels and 66 to 100% reduction in *Lys14* transcript levels, compared to empty vector-transformed controls (Supplementary Figure 8). More specifically, *Lys13* was silenced in all selected plant lines (L3, L11, L14, L17, L26, L27, L47), while only four lines were also silenced for *Lys14* (L11, L17, L27, L47). All *Lys13/14* silenced lines demonstrated statistically significantly lower levels of intraradical FsK colonization when compared to control plants (Figure 5C, p-value = 0.078). Finally, two more available mutant lines, *Ljlys12-3* and the triple mutant *Ljnfr1nfr5lys11*, were included for comparison, since *Lys12* and *Lys11* expression levels were not affected by the presence of the endophyte. The defect in *NF* receptors in the triple mutant was not expected to have an effect on FsK colonization. *Ljlys12-3* displayed significantly higher FsK colonization levels at 4dpi, whereas the triple mutant was colonized to levels similar to those recorded in wt plants (Figure 5B).

## Discussion

### Intraradical accommodation of an endophytic fungus is controlled by multiple LysM receptors

Chitin represents the most ancestral structural polysaccharide in the fungal cell wall (Gow et al., 2017), and most fungi possess chitin synthase enzymes. This was recently, also, shown for oomycetes (Fuechtbauer et al., 2018) (Gibelin-Viala et al., 2019), even though their cell walls primarily consist of β-glucans (Mélida et al., 2013). Chitosaccharides of variable degree of polymerization, produced either by enzymatic cleavage of the long crystalline chitin, or synthesized anew, are considered as Microbe Associated Molecular Patterns (MAMPs), able to trigger symbiotic signalling (as is the case of short chain COs) (Genre et al., 2013), or immune responses (as is the case of longer-chain COs, dp = 6–8 N-acetyl glucosamine) (Stacey and Shibuya, 1997) (Kouzai et al., 2014) (Liang et al., 2014). As regards the receptors of chitinaceous signal molecules from symbionts, highly specific ones have been identified for the NF and Myc LCOs (Amor et al., 2003) (Radutoiu et al., 2003) (Limpens et al., 2003) (Madsen et al., 2003) (Maillet et al., 2011) (Fliegmann et al., 2013) (Murakami et al., 2018). Recent work on structurally related molecules from pathogenic fungi has shown that very similar LysM-RLKs in legumes enable discrimination between chitin perception and perception of NFs (Bozsoki et al., 2017). FsK exudates comprise of, at least partially, chitin-based molecules of unknown degree of polymerization, while heat-labile molecules must also be present. Characterization of the nature of FsK triggering compounds will require substantially additional research but it is notable that heat treatment did not affect the AM fungal exudate activity (Navazio et al., 2007). As anticipated, certain *LysM* genes (*Lys6*, *Lys13*, *Lys14*) are regulated in *Lotus*-FsK interaction, further implying that chitin-based molecules produced by the fungus are recognized by the plant. Two *LysM* genes, *Lys13* and *Lys14* positively control FsK intracellular accommodation and may act as entry receptors of FsK successful passage through the rhizodermis in the legume root. On the contrary, LjLYS6, and perhaps LYS12 may act as balancing receptors, controlling fungal intraradical proliferation, at the post initial infection stages. This role has already been assigned to LYS6 and LYS12 receptors, in interaction of legumes with pathogenic fungi (Bozsoki et al., 2017) (Fuechtbauer et al., 2018).

In the *Lotus*-FsK system, we could detect a phenotype in terms of fungal intraradical accommodation in all available plant lines mutated in *LysM* genes, which are, also, transcriptionally affected by FsK. These RLKs could play a role at both early and late stages of the interaction. Assessing the spatial expression of these genes during FsK progression as well as a putative triggering of immune responses will provide further insight in such roles for the accommodation of the endophytic fungus.

### A very common symbiosis signalling pathway

Structural similarities between chitin-based elicitors from either symbiotic or non-symbiotic microbes raise questions about how plants distinguish friend from foe (Zipfel and Oldroyd, 2017). We investigated the role of the CSSP in establishing an endophytic association, making use of the *Lotus-*FsK system (Skiada et al., 2019). Our results indicate an unparalleled involvement of the CSSP in *Lotus*-FsK association.

Transcriptional reprogramming of CSSP genes, as well as genes acting downstream of it, have been originally described for RL and AM symbiosis. Apart from symbiotic associations in legumes, the involvement of the CSSP in plant associations has been studied in non-symbiotic bacteria in legumes (Sanchez et al., 2005); in legume-parasitic root-knot nematodes (Weerasinghe et al., 2005); and in the successful nodulation of the actinorrhizal plant *Casuarina glauca* by its symbiont, *Frankia sp.*, in which a functional *Lj*SYMRK (Gherbi et al., 2008) as well as a functional *CCaMK* (Svistoonoff et al., 2013) are required. Recently, *L. japonicus* symbiosis genes were shown to structure the root microbiota, as revealed by CSSP mutant analysis (Zgadzaj et al., 2019). In our experimental system, among the genes examined, those acting upstream *CCaMK* had no effect on FsK accommodation in *Lotus* roots and were dispensable for determination of FsK colonization levels. In fact, the non-nodulating and non-mycorrhizal *symRK-1* and *castor-1* mutants (Schauser et al., 1998) (Wegel et al., 1998) (Bonfante et al., 2000) (Novero et al., 2002) (Stracke et al., 2002), displayed a wt-like phenotype in terms of intraradical FsK colonization.

Downstream the nuclear envelope cation channel in the CSSP lies symbiotic calcium spiking (Genre and Russo, 2016). The calcium ion is a universal second messenger in numerous plant signalling pathways. On one hand, cytoplasmic calcium oscillations constitute a general plant cell response to abiotic and biotic stimulation (Knight et al., 1997) (Aldon et al., 2018). More specifically, cytosolic calcium signals have been reported upon perception of symbiotic signals (Navazio et al., 2007), fungal elicitors like chitin and β-glucans (Mithöfer et al., 1999), cell wall fractions and exudates of endophytic fungi (Vadassery et al., 2009) (Johnson et al., 2018). On the other hand, the generation of nuclear and perinuclear calcium oscillations have been shown to trigger specific plant responses. They occur upon abiotic stimulation (Lachaud et al., 2010) (van der Luit et al., 1999), rhizobial and AM fungi perception (Ehrhardt et al., 1996) (Chabaud et al., 2011), perception of Frankia signals by actinorrhizal plants (Chabaud et al., 2016), and perception of flagellin (flg22), oligosaccharides and proteins leading to necrosis in *Nicotiana* sp cells (Lecourieux et al., 2005). Our results show that FsK exudates trigger periodic calcium oscillations in the nucleus of *M. truncatula* ROCs harbouring nuclear-targeted cameleon reporters (Sieberer et al., 2009). The suppression of this response in *dmi2-2* and its persistence in *dmi3-1* mutants, strongly suggests that FsK-triggered calcium signals are acting within the CSSP, in analogy to what has been observed for the response to AM fungi and rhizobia (Wais et al., 2000) (Chabaud et al., 2011) (Genre et al., 2013). To the best of our knowledge, there are no previous reports of nuclear calcium oscillations upon compatible, non-symbiotic legume-microbe interactions that are CSSP-dependent. *M. truncatula* epidermal cells respond to oomycete cell wall fractions by nuclear calcium oscillations but only in a CSSP-independent manner (Nars et al., 2013). This unprecedented conclusion opens a broad field of discussion about the role of the CSSP outside the restricted group of symbiotic interactions *sensu stricto*.

Indeed, a comparison of nuclear calcium oscillations as a response to FsK exudate with calcium signatures in response to symbiotic factors revealed possible analogies: the patterns recorded in our study resemble those triggered by CO4/AM fungal exudate for their relatively irregular peak distribution. In addition, the average peak width of ~60 sec recorded for FsK, is closer to the 71-100 sec interval for significant peaks as response to Nod factors than to the much shorter 24-36 sec interval observed for mycorrhizal factors (Kosuta et al., 2008). Furthermore, by employing the exudates from two additional fungal species as external controls - the pathogenic *Fusarium oxysporum* f. sp. *medicaginis* and the non-mycorrhizal mutualist *P. indica -* we observed comparable nuclear calcium spiking profiles that are CSSP-dependent.

Generation of nuclear calcium oscillations by multiple fungal strains should not be a surprise. Chitin-based molecules are secreted and/or deposited by fungi at the proximity of the interacting plant cell surface. These molecules are structurally related to rhizobial LCOs, as well as Myc COs/LCOs, which are able to activate nuclear calcium spiking in host plants (Ehrhardt et al., 1996) (Sieberer et al., 2009) (Chabaud et al., 2011). Reported symbiotic calcium spiking inducing signals are therefore primarily chitinaceous and the only currently known exception, to our knowledge, is the case of *Frankia* symbiotic factors able to elicit *NIN* (Nodule Inception) gene activation and nuclear calcium spiking on roots of the actinorrhizal plant *Casuarina glauca*. *Frankia* factors are composed of hydrophilic compounds that are resistant to chitinase digestion (Chabaud et al., 2016).

The placement of nuclear calcium spiking at the core of *Lotus*-FsK signalling pathway(s) is supported by CSSP gene expression and phenotyping of mutants impaired in genes acting downstream the nuclear calcium spiking response. *LjCCaMK* (*MtDMI3*), the gene encoding the kinase responsible for deciphering nuclear calcium signals (Lévy et al., 2004), was marginally upregulated both at early and relatively late time points of the interaction. *CCaMK* activation at very early stages of the interaction is of interest, because this gene acts at the core of the signal transduction process. Induction of *CCaMK* at late stages of the interaction, is perhaps associated to fungal progression within host tissues at these stages. Interestingly, *LjCCaMK* expression is only marginally affected by *M. loti* inoculation (Tirichine et al., 2006), suggesting that this calcium-activated enzyme is differentially involved in the two interactions.

An impaired phenotype with lower colonization levels was recorded in both *ccamk-1* and *cyclops-1* mutants, genes acting downstream the calcium spiking response. Interestingly, a comparable delay in nodulation and the occasional and late colonization by AMF (delayed and reduced arbuscule formation) has been reported for the *ccamk-1* weak allele (Schauser et al., 1998) (Demchenko et al., 2004), suggestive of a partial functioning of the truncated protein. Alternatively, at these later time points, and possibly due to fungal overload, the necessity for signal deciphering through CCaMK, and signal transduction through CYCLOPS, may be bypassed by redundant proteins acting in parallel and/or alternative signalling routes that allow the progression of fungal accommodation even in the absence of key components of the CSSP. In this latter case, colonization proceeds fast and reaches wt levels belatedly.

Summarizing the above, we propose that FsK produces at least two different types of molecules to activate the CSSP: chitosaccharides of unknown degree of polymerization and heat sensitive molecules. FsK colonizes the root of legumes at least via three alternative routes: 1) When all CSSP components are functional, FsK utilizes the pathway by generating nuclear calcium spiking and activating CCaMK, which lies at the core of FsK recognition towards intraradical accommodation; 2) When *SYMRK* is impaired, calcium spiking does not occur, but FsK is still capable of effectively colonizing the root, possibly by activating CCaMK through a SYMRK- and calcium spiking-independent route.3) When the calcium spiking response is generated but cannot be perceived or transmitted (impaired *CCaMK* or *CYCLOPS*) the CSSP is completely abolished, and yet FsK is still able to colonize the legume root, though with less efficiency. We assume that in this case, an unexplored third route allows a belated intraradical fungal accommodation.

By moving ‘deeper’ into symbiotic signalling, the nodulation- and mycorrhization-specific transcription factors acting downstream the calcium spiking response, *LjNSP1* and *NSP2* (Maillet et al., 2011) (Delaux et al., 2013) were not affected in our system. Nevertheless, CSSP-regulated genes that act early in nodulation process were induced upon FsK interaction. The constant mild induction of *ENOD40* from 4dpi onwards suggest a possible implication of this nodulin gene in FsK accommodation process. *ENOD40* is an early nodulin gene induced in legume tissues upon NF/chitin pentamer treatment (Minami et al., 1996), during AM infection (van Rhijn et al., 1997) and it is linked to symbiotic (nodule formation) and non-symbiotic organogenetic processes (i.e. lateral root formation) in legumes (Papadopoulou et al., 1996). *ENOD40* is also regulated by cytokinin (van Rhijn et al., 1997). One can speculate that the rapid and sustained expression of *ENOD40* is linked to the preparation of root tissues for FsK accommodation: as a filamentous fungus that grows rapidly within plant tissues, FsK intracellular growth varies from partial to full occupation of the host cell-, involving the rearrangement of plant membranous materials and vesicular activity at infection sites and possibly the assembly of a perifungal membrane system that preserves host cell integrity for a limited time (Skiada et al., 2019). In this frame, plant cells may have to expand, and perhaps divide. Cortical cell division was also recently linked to AM symbiosis (Russo et al., 2018). Analysing the hormonal balance in FsK-legume system during such initial events is expected to shed light on the establishment of this legume-endophyte association.

## Conclusion

In the present study we have shown that multiple LysM receptors that perceive chitin molecules play a role during FsK recognition by the legume plant, and act either as positive or balancing regulators of fungal progression within the root. We furthermore show that the CSSP, so far known to be triggered upon microbial symbiont/symbiotic factor perception to allow intracellular accommodation in legume roots, is also utilized by fungal legume-interacting microbes, with a primary focus on the endophytic *Fusarium solani* strain K. This conclusion suggests that alternative upstream routes - specific to FsK and possibly other microbes - are involved in the activation of the CSSP central core beside rhizobial, mycorrhizal and actinorrhizal factor receptors. The positive regulation of fungal accommodation may be further controlled at steps downstream the calcium spiking response. Our results contribute to the emerging notion that the CSSP is not a sole feature of symbiotic interactions of legumes. It is intriguing and far more perplexing that the question as to how legumes discriminate between two microbes by activating one single pathway should be extended to other interacting microbes as well.

## Materials and methods

### Plant materials

*L. japonicus* wt plants and the CSSP mutant lines *symrk-1*, *castor-1*, *sym15-1 (ccamk-1), sym6-1* (*cyclops-1*) as well as the *LysM* mutant lines *lys6-1*, *lys12-3* and the triple mutant *nfr1nfr5lys11* were used for phenotypic screening of FsK intraradical abundance. All mutant lines were kindly provided by Associate Prof Simona Radutoiu (Department of Molecular Biology and Genetics - Plant Molecular Biology, Aarhus, Denmark). The plants were chemically scarified and grown in Petri dishes as described previously (Skiada et al., 2019) until seedlings transplantation to magenta boxes.

*L. japonicus* wt plants (ecotype ‘Gifu’) were used for gene expression analysis experiments.

*M. truncatula* ROC lines and the CSSP mutant ROC lines *dmi2-2* and *dmi3-1*, expressing the 35S:NupYC2.1 construct (Sieberer et al., 2009) were obtained previously (Chabaud et al., 2011). An apical segment deriving from each ROC line was routinely transferred to square Petri dishes containing M medium (Bécard and Fortin, 1988) and placed in a vertical position at 26°C in the dark, to favour the development of a regular, fishbone-shaped root system (Chabaud et al., 2002). Segments derived from ROCs were used for calcium spiking bioassays upon fungal exudates and chitooligosaccharide treatment.

### Fungal materials

*Fusarium solani* strain K (FsK) (Kavroulakis et al., 2007) was routinely cultured in Potato Dextrose Agar (PDA) medium and conidia were isolated as previously described (Skiada et al., 2019). Fungal conidia were used as inoculum for phenotypic profiling of all mutant lines and for gene expression analysis experiments. *Fusarium oxysporum* f. sp. *medicaginis* (Fom, BPIC 2561) (Snyder and Hansen, 1940) was kindly provided by Benaki Phytopathological Institute (Benaki phytopathological Institute Collection, BPIC), Attiki, Greece. *Piriformospora indica* (MUT00004176) was kindly provided by Mycotheca Universitatis Taurinenesis, Turin, Italy. Fom and *P. indica* were routinely cultured in PDA medium. FsK, Fom and *P. indica* hyphal propagules were used for preparation of fungal exudates for calcium spiking analyses.

### Experimental setup for gene expression analysis and phenotypic screening of *L. japonicus mutants*

*L. japonicus* seedlings (7-11 days old) were transplanted into magenta boxes (3 plants per magenta) containing sterile sand:vermiculite (3:1) and directly inoculated on the root with 10^2^ conidia per plant. Control plants received the same volume of sterile water. The substrate was watered with 30 ml of M medium (Boisson-Dernier et al., 2001) prior to transplantation, in both treatments. Magenta boxes were transferred in a growth chamber (16h light/8h dark photoperiod, 22°C).

For gene expression analysis, plants were harvested at 1, 2, 4, 6, 12 days post inoculation (dpi), washed 5x with sterile water to remove the excess extraradical mycelium, root tissues were collected, flash frozen under liquid nitrogen and kept at −80°C until subsequent DNA/RNA isolation. Four (4) biological replicates were assessed for each treatment, with each replicate consisting of 3 individual plants. Estimation of intraradical abundance and gene expression analysis were performed as described below. For phenotypic profiling, wt and CSSP mutants were harvested at 4 and 8 dpi and *LysM* mutants were harvested at 4 dpi. Different batches of wt *L. japonicus* plants were assessed for CSSP or for *LysM* mutants. Harvested plants were surface sterilized (1% v/v NaOCl), washed 5x with sterile water, root tissues were collected, frozen under liquid nitrogen and kept at −20 until subsequent DNA isolation. Five (5) biological replicates were assessed for each treatment, with each replicate consisting of 3 individual plants. Estimation of intraradical fungal abundance was performed as described below.

### *Agrobacterium rhizogenes*-mediated hairy-root transformation and plant inoculation with FsK

A 176 bp fragment specific of *Lys13*/*Lys14* genes (100% nucleic acid similarity in both genes) was amplified by PCR using specific primers (Supplementary Table 2) and cDNA derived from RNA extracted from *L. japonic*us root tissues as a template, and was introduced in the pUBI-GWS-GFP vector (Maekawa et al., 2008) in order to generate an RNA-hairpin expression construct targeting both *Lys13*/*Lys14*. The binary vector was transformed into *Agrobacterium rhizogenes* LBA1334 and used for generation of *L. japonicus* hairy roots as described previously (Krokida et al., 2013). Control plants were transformed with the empty pUBI-GWS-GFP. Regenerated hairy roots were screened for GFP fluorescence, non-transformed roots excised, and the presence of the hairpin expressing T-DNA was verified by PCR using primers specific for the GFP CDS.

Plants with regenerated hairy roots (16 days old) were transferred into pots containing sterile sand:vermiculite (3:1) substrate, in a growth chamber (16h light/8h dark photoperiod, 22°C). Plants were alternately watered with distilled water or Hoagland medium. Two days post transfer plants were inoculated with 10^2^ FsK conidia per root. At 5 dpi, plants were harvested, washed 5x with sterile water to remove the excess extraradical mycelium, root tissues were collected, flash frozen under liquid nitrogen and kept at −80°C until subsequent DNA/RNA isolation.

Silencing of *Lys13*, *Lys14* was verified by gene expression analysis. RNA isolation, cDNA synthesis and qPCR for estimation of *Lys13/Lys14* transcript levels was conducted as described below. Based on these results 7 lines were chosen showing reduced expression levels of *Lys13*, *Lys14* or both. DNA was isolated from the same tissues and 1μl was used for absolute quantification of *Fusarium solani* ITS gene copy number and therefore estimation of intraradical fungal abundance.

### DNA isolation

*L. japonicus* root tissues were grounded with a pestle, under the presence of liquid nitrogen, and total DNA was isolated using the CTAB method (Doyle and Doyle, 1987). For quantification of intraradical fungal abundance, the weight of the plant tissue was determined prior to DNA isolation. DNA concentration was determined using Qubit 2.0 fluorometer. 1μl of the eluted DNA was used as template in qPCR reaction for estimation of intraradical fungal abundance in *L. japonicus* wt, *CSSP* and *LysM* mutant/RNAi lines.

### RNA isolation

Total RNA was extracted using the Isolate II RNA Plant Kit (BIO-52077, BIOLINE) according to manufacturer’s instructions. To eliminate genomic DNA carry-over in subsequent reactions, samples were treated with DNase I (18047019, Invitrogen) at 37°C for 1h. Elimination of genomic DNA contamination was verified by PCR, using primers specific for *Lj*UBIQUITIN gene (Supplementary Table 2) and subsequent agarose gel electrophoresis of the PCR products.

### Gene expression analysis

First strand cDNA was synthesized from DNase treated total RNA, using the PrimeScript™ 1st strand cDNA Synthesis Kit (6110A, Takara). Quantitative real-time RT-PCR (qPCR) was performed using gene-specific primers (Supplementary Table 1) and the fluorescent intercalating dye Kapa SYBR Green (Kapa SYBR FAST qPCR Master Mix Universal, Kapa Biosystems) in a BIORAD CFX Connect^TM^ Real-Time PCR Detection System. All quantifications were normalized to *Lj UBIQUITIN* housekeeping gene. Data were analysed according to (Pfaffl, 2001). Reaction efficiencies were estimated through the free software program LinRegPCR (Ramakers et al., 2003).

### Quantification of fungal colonization in *L. japonicus* wt, CSSP, *LysM* mutant/RNAi lines

To estimate fungal abundance within root tissues, absolute quantification of *F. solani ITS* or *TEF1a* gene was performed by using a previously constructed standard curve (Skiada et al., 2019). For each experiment, amplification of each sample (standards - unknowns) occurred in a 10 μl reaction mixture containing Kapa SYBR FAST qPCR Master Mix Universal (1x), 200 nM of each primer, and 1 μl of DNA, using a thermocycling protocol of 3 min at 95 °C; 45 cycles of 15 s at 95 °C, 20 s at 58 °C, followed by a melting curve to check the specificity of the products. Absolute quantification of unknown samples was estimated as the exact copy concentration of the target gene by relating the CT value to the standard curve. The fungal gene copy numbers of all samples were normalized by ng of total DNA isolated, or by mg of tissue used to extract DNA, or both. Agarose gel (1.5% w/v) electrophoresis of qPCR products was routinely performed, to verify the amplification of a single fragment of the desired length. Intraradical fungal colonization of *L. japonicus* control/FsK inoculated plants was estimated by quantification of *TEF1a* fungal gene. qPCR amplification efficiencies were 100.91 % with r^2^ value of 0.991 and a slope of −3.300. Intraradical fungal colonization in *L. japonicus* wt-CSSP mutants, in wt-*LysM* mutants and wt-RNAi lines was estimated by quantification of *ITS* fungal gene. qPCR amplification efficiencies were 91.82 % with r^2^ value of 0.997 and a slope of −3.535 for the *CSP* mutants, 95.46 % with r^2^ value of 0.995 and a slope of −3.436 for *LysM* mutants and 89.7 % with r^2^ value of 0.997 and a slope of −3.598 for the RNAi lines screening.

### Preparation of fungal exudates and root treatment

Exudates of FsK, Fom and *P. indica* were prepared as previously described (Lace et al., 2015). Briefly, 1ml of water was distributed in eppendorf tubes and inoculated with fungal hyphae obtained by slightly scrapping with a sterile scalpel the surface of a previous culture of the fungus on Potato Dextrose Agar (PDA). Tubes were incubated for 7 days at 26°C, the exudate was afterwards recovered with a sterile syringe, filter sterilized (0.2 μm filters), concentrated 10-fold using a lyophilizer, and stored at −20°C until further use. Exudates of *Gigaspora margarita* were prepared as described in (Chabaud et al., 2011). Segments of primary *Mt* wild type and *dmi2-2*/*dmi3-1* mutant ROCs carrying one or two young laterals were placed in a microchamber and treated with 100 μl of fungal exudate as described in (Chabaud et al., 2011). For CO5 treatment, aqueous solutions at a final concentration of 10^−6^ M were prepared from a concentrated stock of purified chitin pentamer solution and root explants were treated as above. As negative control, sterile water was used. As positive control, *G. margarita* germinated spore exudate, or purified COs (degree of polymerization, n=5) were used.

### Assays performed on fungal exudates and COs

Chitinase treatment of the *G. margarita* GSE was performed using 1 mg ml^−1^ chitinase from *Streptomyces griseus* (ref C6137; Sigma-Aldrich) in sterile H_2_O for 16 h at room temperature. Autoclave treatment was performed by autoclaving the exudates (121 °C, 1 atm) for 20 min or for 1 h. Chitinase treated exudates of FsK and Fom were tested for their ability to generate calcium spiking responses on wt NupYC2.1 *Mt* ROCs epidermis. Autoclave treated exudates of FsK were tested for their ability to generate calcium spiking responses on wt NupYC2.1 *Mt* ROCs epidermis.

### Confocal microscopy for calcium spiking measurements in *M. truncatula* ROC epidermis

A FRET-based ratiometric approach (Sieberer et al., 2009) was used to record the relative changes of nuclear calcium concentrations corresponding to Yellow Fluorescent Protein (YFP) to Cyan Fluorescent Protein (CFP) fluorescence intensity changes over time (Miyawaki et al., 1997). Analysis was performed using a Leica TCS SP2 confocal microscope fitted with a long distance x40 water-immersion objective (HCX Apo 0.80). Measurements were performed and settings were set as previously described (Chabaud et al., 2011). The pinhole diameter was set at 6 airy units. YFP and CFP fluorescence intensities were calculated for each region of interest (each nucleus) using Leica LCS software. Values were then exported to a Microsoft Excel spreadsheet, the YFP/CFP ratio for each time frame was calculated, and ratio values were plotted over time. A graphical representation of FRET ratio was finally generated, which indicated the relative calcium concentration changes in the nucleus over time. Nuclear spiking was examined in all cases in an atrichoblast/low-trichoblast zone located 10-20 mm from the root tip.

Calcium spiking measurements were furthermore analyzed using CaSA software (Russo et al., 2013). The generation of at least 3 peaks was the threshold for discrimination between responding and non-responding cells. The total number of epidermal cell nuclei and independent root segments analyzed for each test are reported in the corresponding Figures/Figure legends. CaSA output PNG image files indicating spike initiation and end points, were used to manually calculate peak width and lag phases prior to calcium spiking response initiation in responding nuclei. CaSa output waiting time autocorrelation values were used to generate the corresponding histogram.

### Statistical analysis

Two-way ANOVA followed by Tukey’s post-hoc test was used in gene expression analysis data. Student’s t-test was used in all pairwise comparisons. One-way ANOVA was used in comparisons between 3 or more independent (unrelated) groups (i.e. analysis of calcium spiking results obtained after plant cells elicitation with the three investigated fungal exudates). The non-parametric Kolmogorov-Smirnov (K-S) two-sample test was used to assess whether data corresponding to a) number of peaks, b) lag phase prior to spiking initiation, c) width of peaks, generated in *Mt* nuclei by the 3 elicitors, are drawn from the same distribution (pairwise comparisons among results from the 3 elicitors). Pearson’s chi squared test was used to test whether the frequency distribution of waiting times autocorrelation values of peaks generated by the 3 fungal exudates are independent. Statistical analyses used are indicated in detail in the corresponding Figures/Figure legends.

## Acknowledgement

We are grateful to Daniela Tsikou for sharing ideas and her suggestions during the initiation of the project, and for reading and commenting on the manuscript; to Constantine Garagounis for his suggestions and contribution in the analysis of hairy roots experiments, and for reading and commenting on the manuscript. We also thank Dimitra Papantoniou for her assistance in the hairy root experiment. This work was partially supported by the Postgraduate Programme 3817 of the Department of Biochemistry and Biotechnology, University of Thessaly, Programme 4757.17 (funded by GGSRT/EU/ NSRF to KKP) and COST Αction FA1206 and FA1405 (STSM Grants to VS).

