## Supplemental files for "Symbiotic signalling is at the core of an endophytic *Fusarium solani*-legume association"

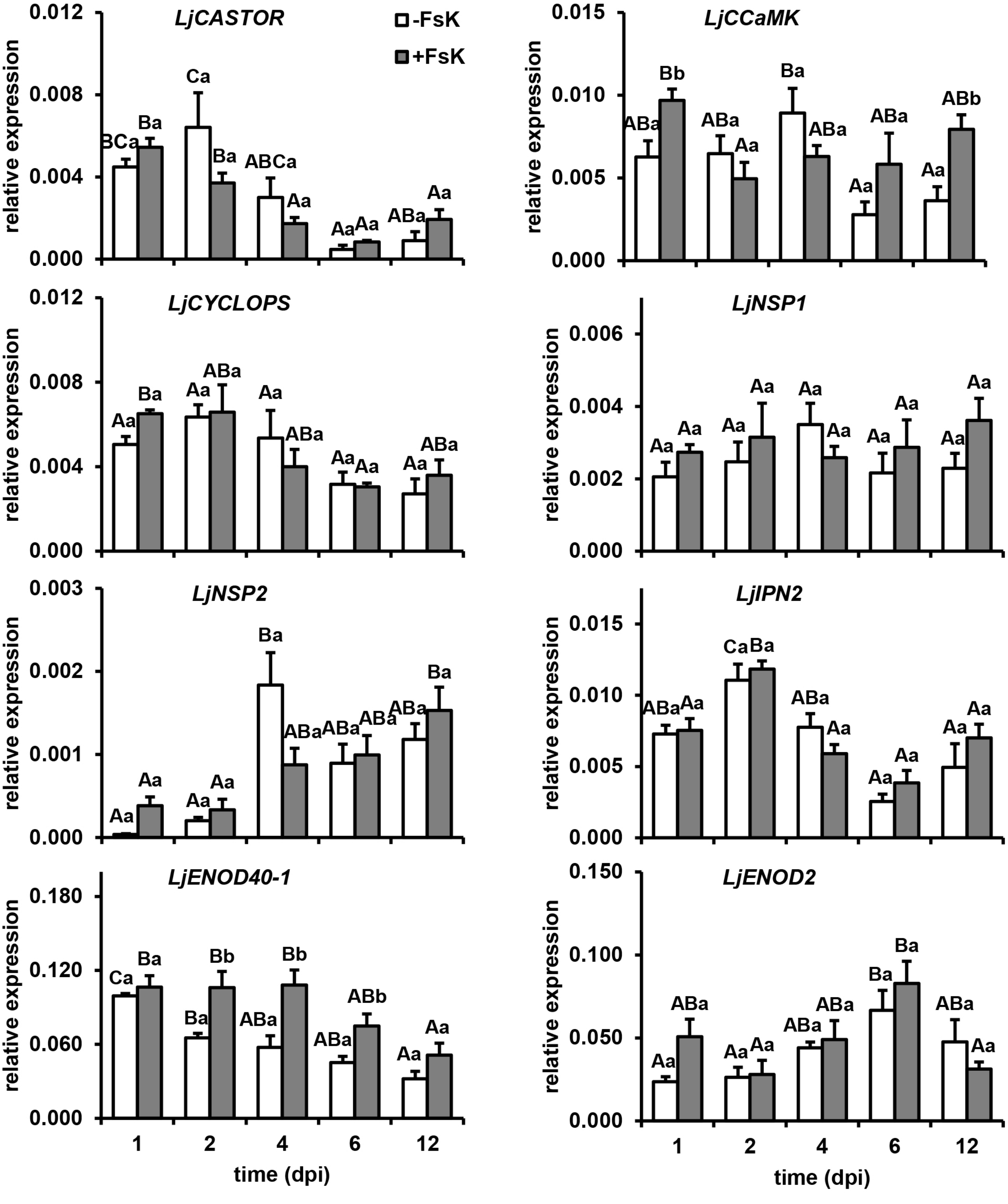


**Supplementary Figure 1 Relative expression of *L. japonicus* genes acting within and downstream the CSSP, upon FsK treatment.**

Temporal quantitative gene expression analysis of *L. japonicus* CSSP genes: *CASTOR*, *CCaMK*, *CYCLOPS*, and genes acting downstream the CSSP: *NSP1*, *NSP2*, *IPN2*, *ENOD40-1* and *ENOD2*, in control and FsK-inoculated *L. japonicus* root tissues. Expression levels are normalized to *Lj UBIQUITIN* housekeeping gene. Data are presented as means and standard errors of 3-4 biological replicates of 3 plants.

Different upper case letters denote significant differences within the same treatment at the 0.05 level, whereas different lower case letters, significant differences within the same time at the 0.05 level (two-Way Anova, Tukey's test).

White/grey bars, expression in control/FsK inoculated tissues

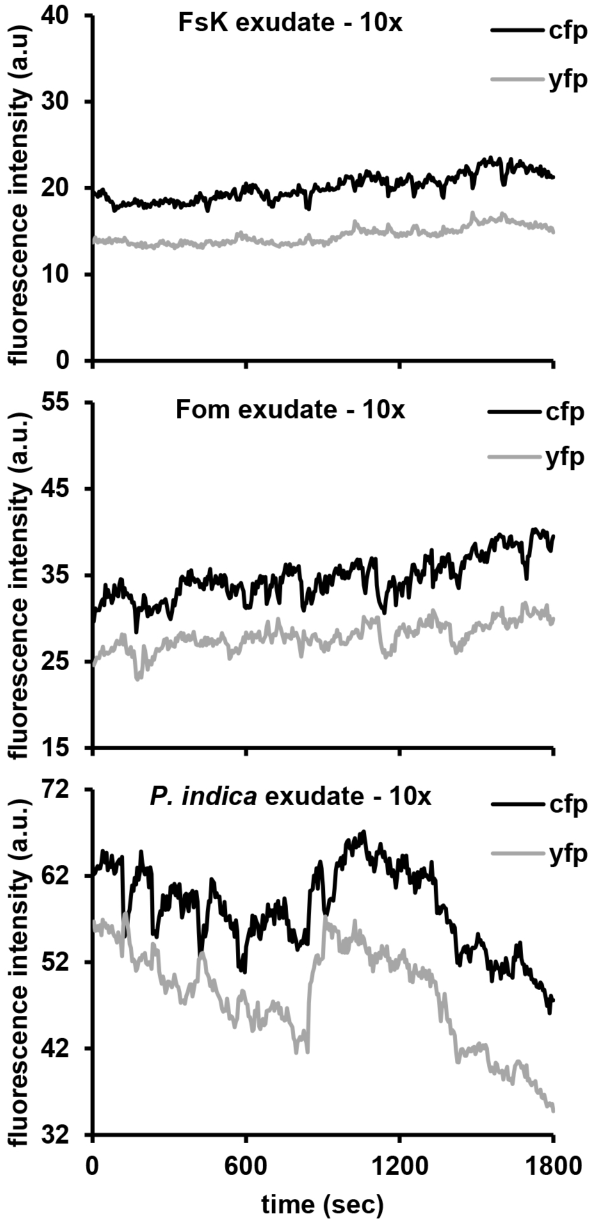


**Supplementary Figure 2 Changes in CFP, YFP fluorescence over time obtained from *Mt* wt ROC epidermal cells upon elicitation with fungal exudates.**

Calcium spiking responses were recorded in a low-trichoblast-region located 10-20 mm from the root tip. *M. truncatula* wt ROCs expressing the NupYC2.1 cameleon were used for the bioassay. Fungal exudates derived from FsK, Fom, or *P. indica* were used as elicitors.

Graphs show the changes in CFP and YFP fluorescence (in arbitrary units) following addition of FsK, Fom or *P. indica* exudate, respectively. The presented plotted changes in fluorescence correspond to YFP:CFP ratios presented in Figure 2.

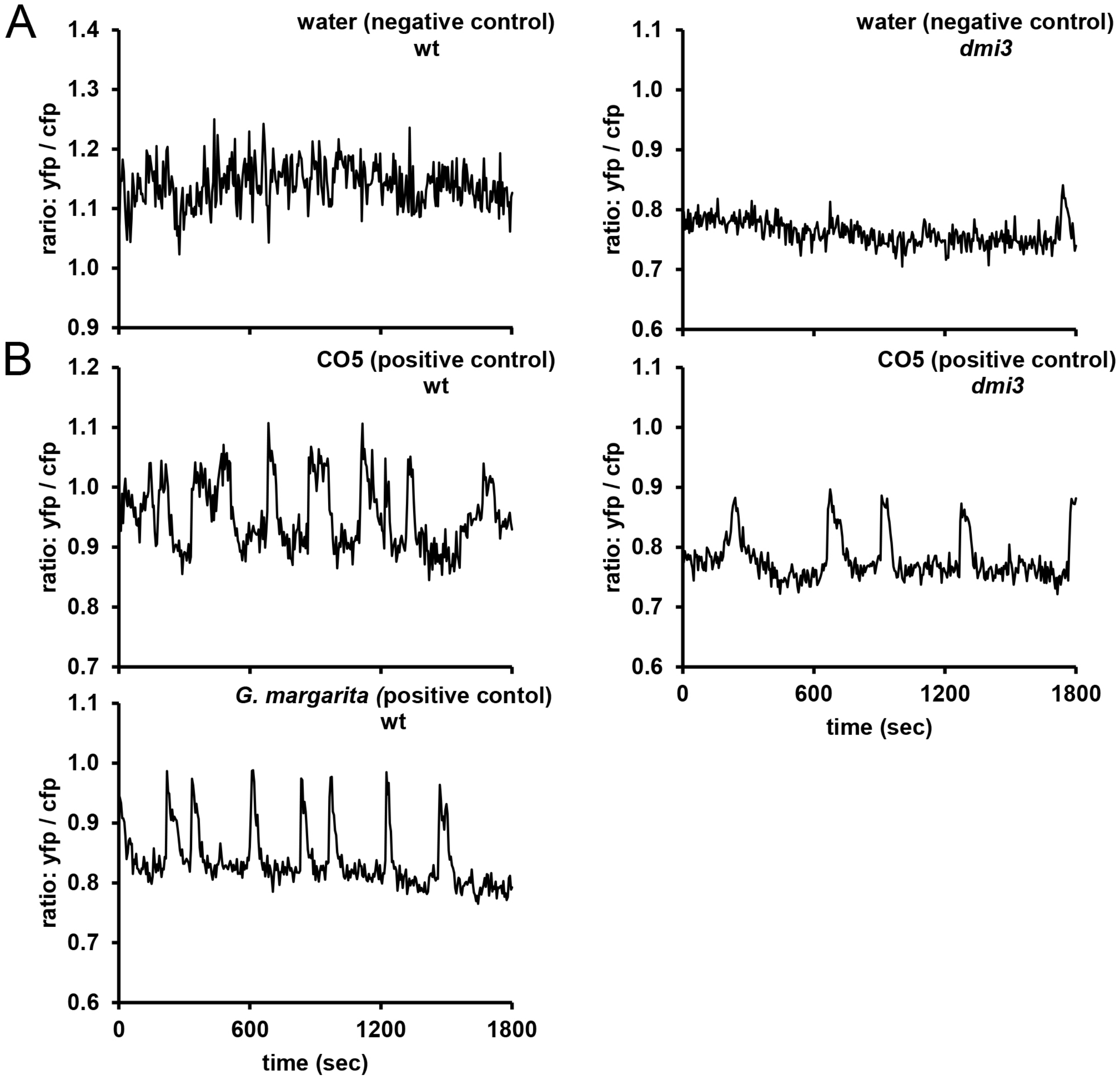


**Supplementary Figure 3 Absence of calcium spiking in response to mock treatment, and generation of calcium spiking in response to CO5 and *G. margarita* germinated spore exudate treatment.**

Calcium spiking responses were recorded in a low-trichoblast-region located 10-20 mm from the root tip. *M. truncatula* wt and *dmi3-1* ROCs expressing the NupYC2.1 cameleon were used for the bioassay.

**A - B** Graphs show ratios of YFP:CFP fluorescence over time obtained from epidermal cells.

**A** Plots showing absence of changes in nuclear calcium levels in wt or *dmi3-1* ROC lines upon (mock) treatment with sterile water.

**B** Plots showing nuclear calcium spiking induced in wt ROC lines after treatment with chito-oligosaccharides (CO5) or *G. margarita* germinated spore exudate, and in *dmi3-1* ROC lines after treatment with CO5.

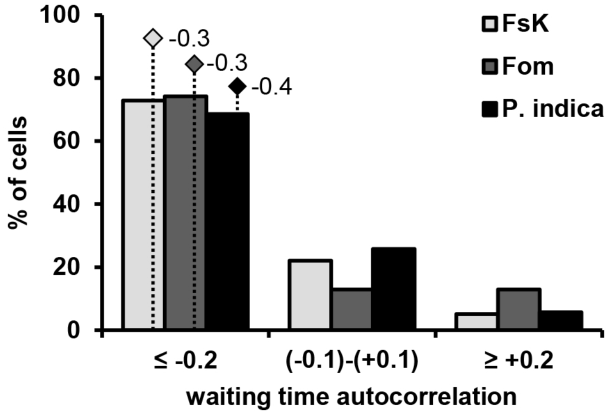


**Supplementary Figure 4 Distribution of waiting time autocorrelation values in the spiking profiles induced by fungal exudates.**

Histogram illustrating the waiting time autocorrelation values in the spiking profiles induced by FsK, Fom, or *P. indica* fungal exudate. The majority of nuclei demonstrate negative autocorrelation values (≤ -0.2) in all fungal exudates tested.

Statistical analysis showed no dependence between the fungal exudate tested and the frequency distribution of waiting time autocorrelation values, at the 0.05 level (Pearson's chi-squared test).

Light grey/dark grey/black bars: frequency of waiting time autocorrelation values intervals in response to FsK/FoM/*P. indica* exudate, respectively.

Waiting time autocorrelation values correspond to peaks derived from active plant cell nuclei of biological replicates, as presented in Supplementary Table 1.

Rhomboid shapes, waiting time autocorrelation median values of peaks generated in response to FsK/FoM/*P. indica* exudate, respectively.

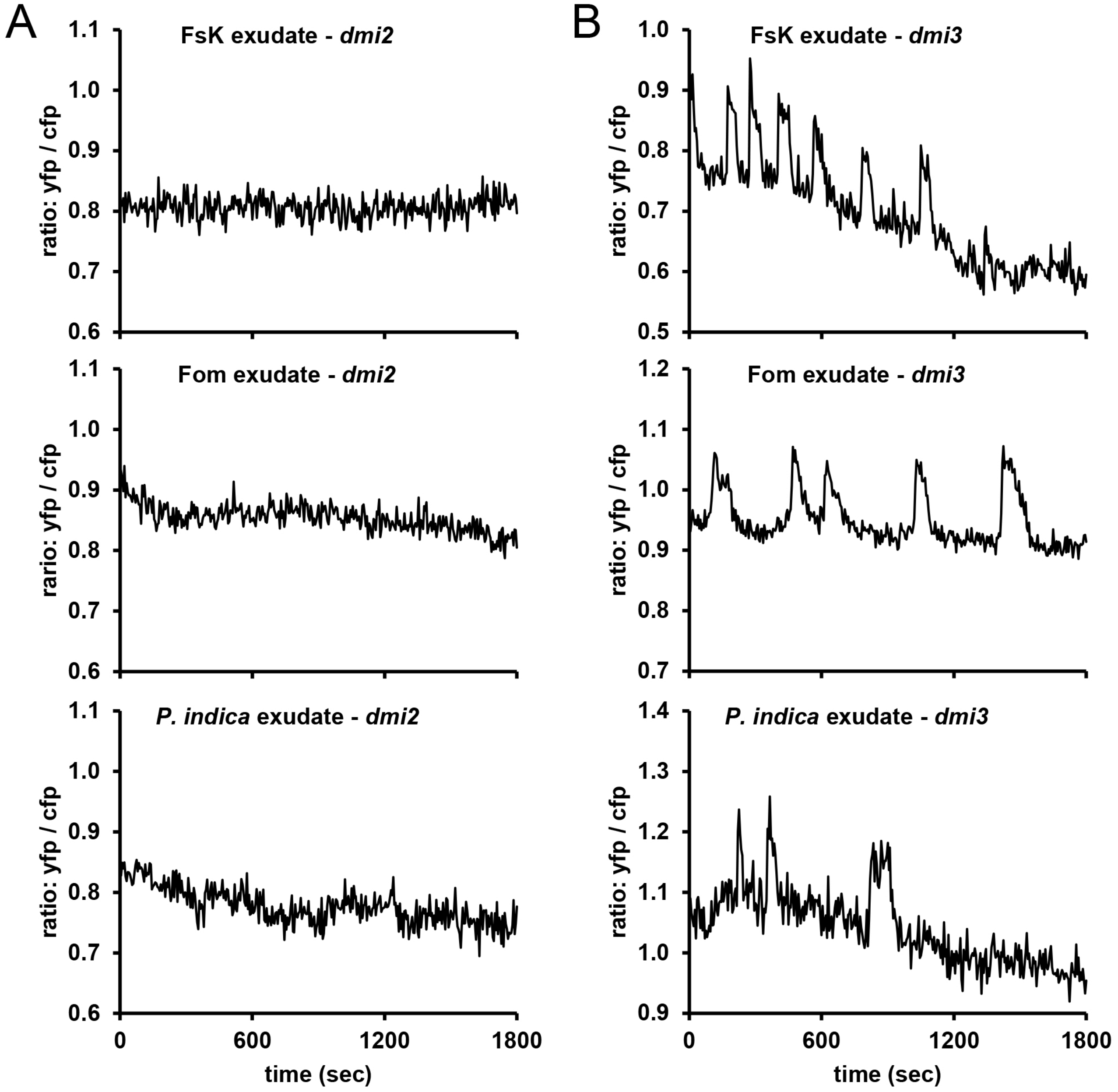


**Supplementary Figure 5 Nuclear calcium spiking is abolished in the *Mt dmi2-2*, but maintained in the *Mt dmi3-1* mutant lines.**

Calcium spiking responses were recorded in a low-trichoblast-region located 10-20 mm from the root tip. *M. truncatula dmi2-2* and *dmi3-1* ROCs expressing the NupYC2.1 cameleon were used for the bioassay. Fungal exudates derived from FsK, FoM, or *P. indica* were used as elicitors.

**A - B** Graphs show ratios of YFP:CFP fluorescence obtained from epidermal cells in a low-trichoblasts-zone.

**A** Representative plots showing absence of changes in calcium levels in the nucleus of *dmi2-2* lines post treatment with 10x concentrated FsK, FoM, or *P. indica* exudate.

**B** Representative plots showing the induced changes in calcium levels in the nucleus of *dmi3-1* lines post treatment with 10x concentrated FsK, FoM, or *P. indica* exudate. All graphs show the plotted changes in the ratio of YFP:CFP fluorescence.

S6

A

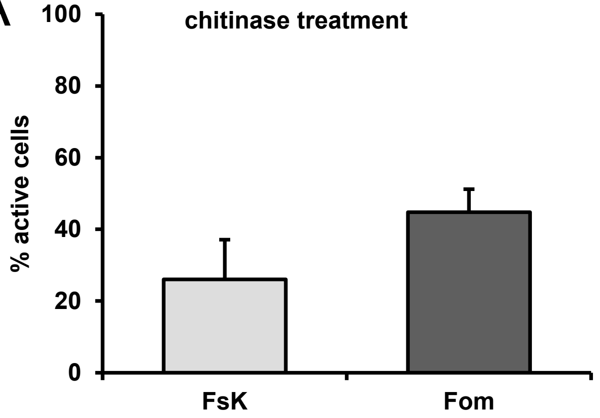


B

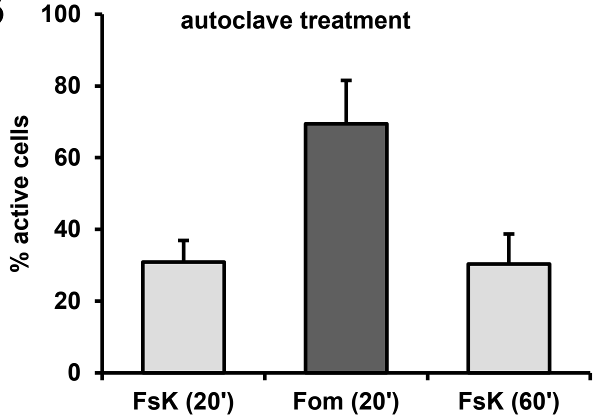


C

| treatment | fungal exudate | ROC lines | ROC segments (biological replicates) | nuclei screened |
|---|---|---|---|---|
| chitinase | FsK | wt | 4 | 45 |
| | Fom | wt | 3 | 35 |
| autoclave (20 min) | FsK | wt | 5 | 64 |
| | Fom | wt | 4 | 34 |
| autoclave (1 h) | FsK | wt | 3 | 41 |

D

| Student's t-test | | | |
|---|---|---|---|
| time of treatment | groups compared | p-value | |
| 16h | FsK NT - FsK CT | 0.02 | * |
| | Fom NT - Fom CT | 0.02 | * |
| 20 min | FsK NT - FsK ACL | 0.01 | * |
| | Fom NT - Fom ACL | 0.46 | *ns* |
| 1 h | FsK NT - FsK ACL | 0.02 | * |

**Supplementary Figure 6 Nuclear calcium spiking responses in *Mt* wt ROCs elicited by FsK/FoM fungal exudate pretreated with chitinase or heat.**

Calcium spiking responses were recorded in a low-trichoblast-region located 10-20 mm from the root tip. *M. truncatula* wt ROCs expressing the NupYC2.1 cameleon were used for the bioassay. Fungal exudates derived from FsK or Fom were used as elicitors.

**A** Chitinase pretreated FsK or FoM exudate elicits nuclear calcium spiking in a statistically significant lower number of cells.

**B** Heat treatment of the fungal exudates, elicits nuclear calcium spiking in a statistically significant lower number of cells only in the FsK fungal exudate. Extending the period of heat treatment from 20 min to 1 h does not further affect the number of cells showing calcium spiking upon treatment with FsK fungal exudate.

Light/dark grey bars: percentages of cells responding with calcium spiking to FsK/Fom treated exudate, respectively.

Data for panel A and B are presented as means and standard errors of biological replicates (individual lateral root segments derived from ROCs) as indicated in panel C.

**C** Summary of ROC segments (biological replicates), and total nuclei screened upon application of the treated with chitinase or heat FsK/FoM fungal exudate.

**D** Summary of p-values estimated based on Student's t-test (0.05 level). Comparisons were conducted in % of cells responding to untreated versus treated fungal exudates.

% of cells responding to untreated fungal exudates are shown in Figure 3A.

NT, non-treated exudate; CT, chitinase treatment; ACL, autoclave treatment

S7

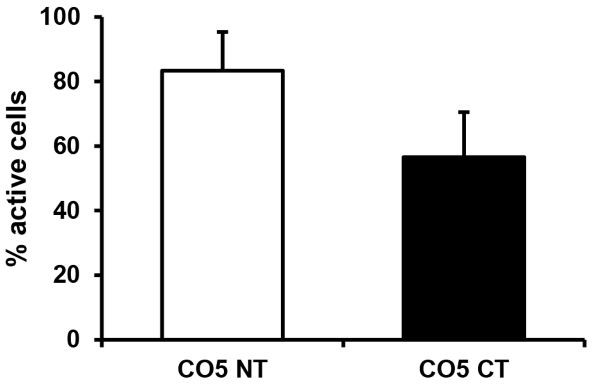


**Supplementary Figure 7 Partial abolishment of nuclear calcium spiking in *Mt* wt ROC epidermis upon chitinase treatment of chitooligosaccharides.**

Calcium spiking responses were recorded in a low-trichoblast-region located 10-20 mm from the root tip. *M. truncatula* wt ROCs expressing the NupYC2.1 cameleon were used for the bioassay. Untreated CO5 solution and CO5 solution enzymatically treated with *S. griseus* chitinase were used as elicitors. Percentage of responsive cells (peak number>2), in 30 min recordings post CO5 (non-treated and chitinase-treated) elicitation.

Data are presented as means and standard errors of 3 biological replicates (individual lateral root segments derived from ROCs).

Statistical analysis showed no statistically significant difference in the % of cells responding to non-treated versus chitinase-treated CO5 solution at the 0.05 level (Student's t-test).

White/black bars: % of responsive cells upon non-treated (CO5 NT) or chitinase-treated (CO5 CT) elicitation, respectively.

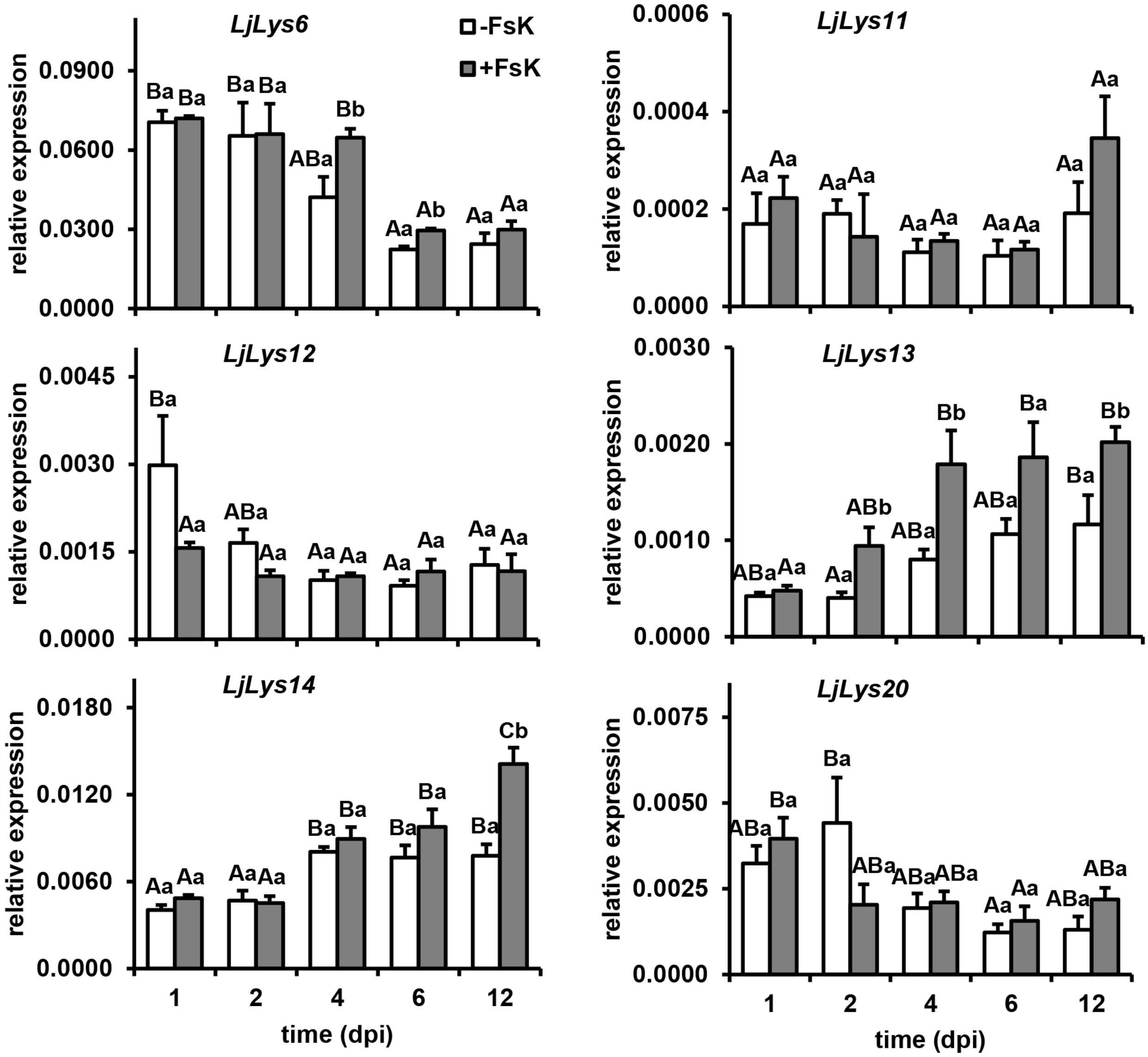


**Supplementary Figure 8 Temporal gene expression analysis of *L. japonicus LysM* genes upon FsK treatment.**

Temporal quantitative RT-PCR analysis of *Lj Lys6*, *Lys11*, *Lys12*, *Lys13*, *Lys14*, and *Lys20* transcripts in control and FsK-inoculated *L. japonicus* root tissues. Expression levels are normalized to *Lj UBIQUITIN* housekeeping gene. Data are presented as means and standard errors of 3-4 biological replicates of 3-4 plants.

Different uppercase letters denote significant differences within the same treatment at the 0.05 level, whereas different lowercase letters, significant differences within the same time at the 0.05 level (two-Way Anova, Tukey's test).

White/grey bars, expression in control/FsK inoculated tissues, respectively.

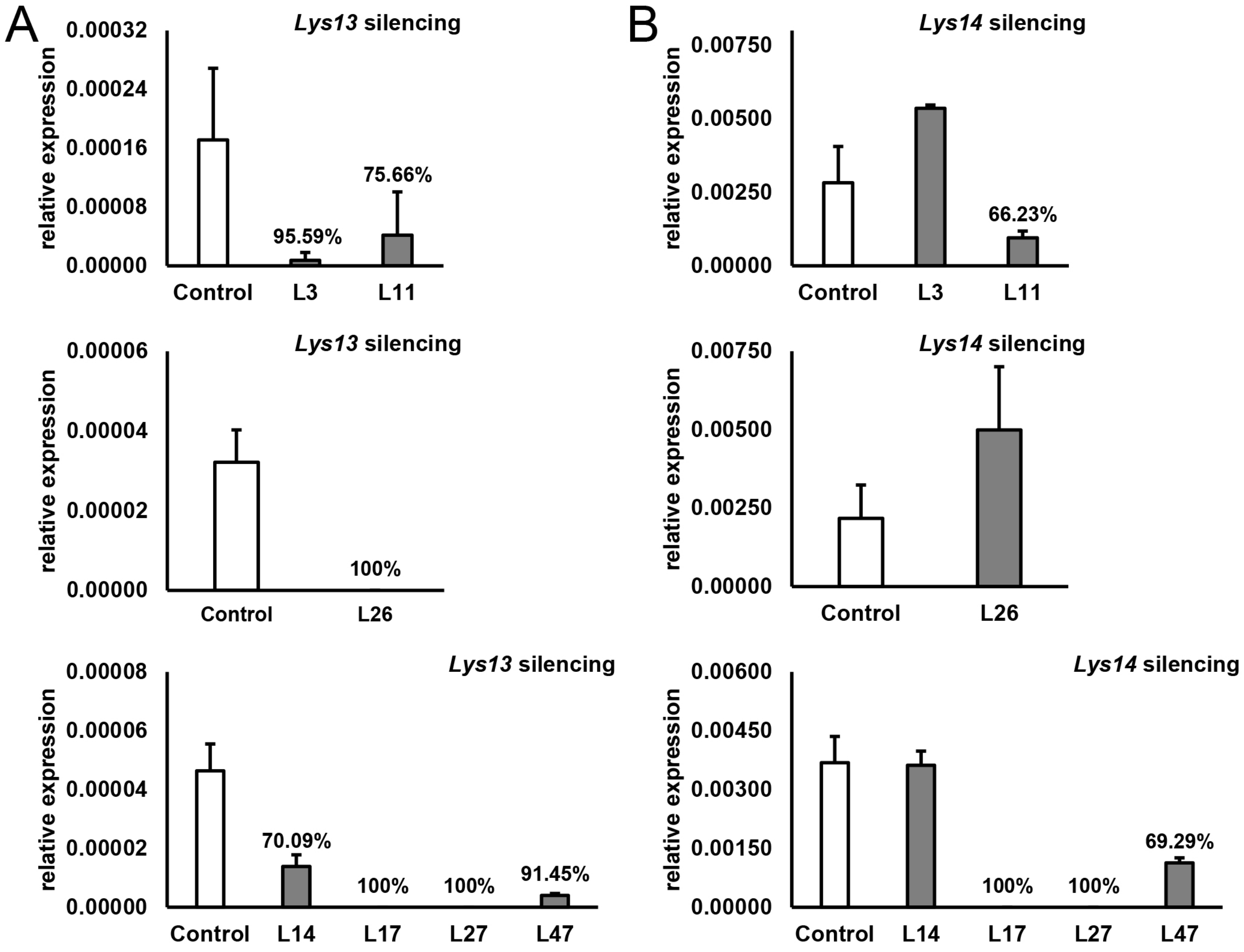


**Supplementary Figure 9 Verification of *Lys13* and/or *Lys14* gene silencing in *Lys13/14* RNAi lines**

Gene expression analysis of *Lys13* and *Lys14* gene in control (transformed with the empty vector) and *Lys13/14* RNAi lines. 7 lines were selected and presented, based on their lower *Lys13* (A) and/or *Lys14* (B) relative transcript levels. The % reduction in each gene relative expression is presented above each respective bar.

Data for control lines are presented as means and standard errors of 3-4 biological replicates.

Data for control corresponding to line 26 (second graph for both panel A and B) are presented as mean values and standard deviation of 2 technical replicates of one single biological replicate.

Data for RNAi lines are presented as mean values and standard deviation of 2 technical replicates.

**S10**

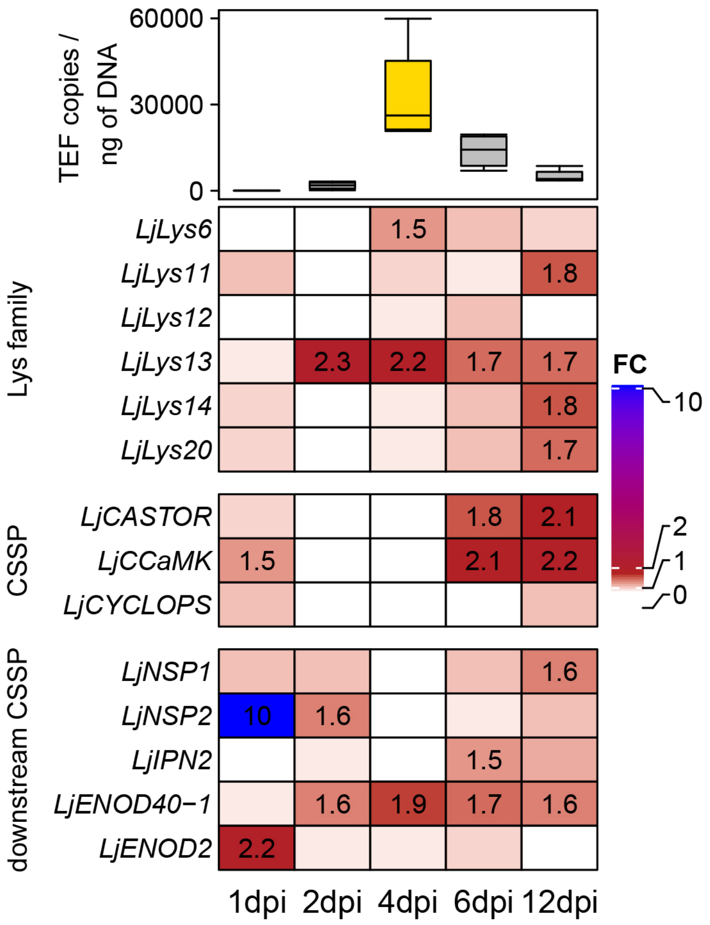


**Supplementary Figure 10 Heatmap summarizing fold-changes in relative transcript levels of genes examined in the present study**

Temporal fold-changes in transcript levels of all genes investigated in the present study are summarized and presented as a heatmap. Fold-change values are indicated within each cell, only FC>1.45 are presented. Gene expression analysis is also presented in detail in Supplementary Figures 1 and 8.

Colour code: 0<FC<1, white; 1<FC<2, firebrick; 2<FC<11, blue.

Top annotation of the heatmap: Box-plots represent the temporal quantification of fungal abundance within FsK-inoculated *L. japonicus* roots tissues, via qPCR, using primers specific for a fragment of *TEF1a* gene. Tissues were collected at 1, 2, 4, 6, 12 dpi. Values are normalized to ng of total DNA isolated.

Boxplots represent the values of 4 biological replicates per time point; each biological replicate consisting of 3 plants.

Different box colours indicate significant differences in fungal abundance (gene copy number average values), among the different time points investigated, at the 0.05 level (one-way Anova, Tukey's test).

ST1

| **A. Total nuclei screened upon elicitation of *Mt* wt ROC lines with fungal exudates** | | | |
|---|---|---|---|
| **Fungal exudate** | **ROC lines** | **ROC segments (biological replicates)** | **Nuclei screened** |
| FsK | wt | 7 | 85 |
| Fom | wt | 3 | 38 |
| *P. indica* | wt | 6 | 78 |
| **B. Total nuclei screened for calculations of delay time and total peak number screened for calculations of peak width** | | | |
| **Fungal exudate** | **ROC lines** | **Nuclei screened for initial delay time** | **Peaks screened for width** |
| FsK | wt | 59 | 316 |
| Fom | wt | 31 | 156 |
| *P. indica* | wt | 34 | 135 |
| **C. Total nuclei screened upon elicitation of *Mt dmi2-2* and *dmi3-1* with fungal exudates** | | | |
| **Fungal exudate** | **ROC lines** | **ROC segments (biological replicates)** | **Nuclei screened** |
| FsK | *dmi2-2* | 3 | 42 |
| | *dmi3-1* | 6 | 70 |
| Fom | *dmi2-2* | 3 | 28 |
| | *dmi3-1* | 4 | 43 |
| *P. indica* | *dmi2-2* | 3 | 37 |
| | *dmi3-1* | 3 | 36 |

**Supplementary Table 1 Summary of individual *Mt* ROC segments (biological replicates), total plant cell nuclei, and total number of peaks screened for the measurements presented.**

**A** Summary of individual ROC segments (biological replicates), and total plant cell nuclei screened upon application of fungal exudates

**B** Summary of total plant cell nuclei used to calculate delay time, and total peaks used to calculate spike width upon application of fungal exudates. Only responsive cells (peak number>2) were used for the analysis.

**C** Summary of the total number of *dmi2-2* and *dmi3-1* ROC segments (biological replicates), and total nuclei screened upon FsK/FoM/*P. indica* exudate treatment. The total number of cells (active and non-active) was used in the analysis and is presented.

**Supplementary Table 2. Primer sequences used in the present work**

| Target | Sequence (5'-3'), (F, forward; R, reverse) | Reference |
|---|---|---|
| **Check for genomic DNA contamination in DNase treated RNA** | | |
| *LjUBIQUITIN* | F: ATGCAGATCTTTTGTGAAGAC | (Flemetakis et al., 2000) |
| | R: ACCACCACGGAAGACGGAG | (Flemetakis et al., 2000) |
| **Expression analysis (qPCR)** | | |
| *LjUBIQUITIN* | F: TTCACCTTGTGCTCCGTCTTC | (Hayashi et al., 2005) |
| | R: CCAGAAGAGGCCACAACAAAC | |
| *LjLys6* | F: TTCAGTTCGCAAGATGGCCC | (Lohmann et al., 2010) |
| | R: AACATCCCAGTCGTCAGTGG | (Lohmann et al., 2010) |
| *LjLys11* | F: AGCCTCTGTCAAGACCAACC | (Lohmann et al., 2010) |
| | R: CCCACAAGTCCATGATCTCTC | (Lohmann et al., 2010) |
| *LjLys12* | F: ATGGACCCTTGAGTGAGTGG | (Lohmann et al., 2010) |
| | R: TGGATGTGAGGAGGAGAAGTG | (Lohmann et al., 2010) |
| *LjLys13* | F: GTTGAACGCTCCAGATCTGTTAGC | (Lohmann et al., 2010) |
| | R: GTGTAGTGAGGGTGAAAATGATCC | (Lohmann et al., 2010) |
| *LjLys14* | F: GGGATCCATCTGATGAACTC | (Lohmann et al., 2010) |
| | R: CACAGAATTTATTATGGCTTCAG | (Lohmann et al., 2010) |
| *LjLys20* | F: CCACATCCTTCTAAGTACATGTTAGG | (Lohmann et al., 2010) |
| | R: ACCATGATTATTGCGCCGCCAG | (Lohmann et al., 2010) |
| *LjCASTOR* | F: ATGGTGGCCTTGACATAAG | (Imaizumi-Anraku et al., 2005) |
| | R: AGTGACGACGTATAACAGCA | (Imaizumi-Anraku et al., 2005) |
| *LjCCaMK* | F: TCCCTCTTAGCAACCGCGTATA | |
| | R: AGCACAAACGGAGAGCATCA | |
| *LjCYCLOPS* | F: GCTGGCAGATGAAAAAGAGC | (Yano et al., 2008) |
| | R: GCGTGTTTGAGCACAACATT | (Yano et al., 2008) |
| *LjNSP1* | F: ATTCCCATCCAGCTTCCAC | (Lopez-Gomez et al., 2012) |
| | R: GAGGTCGAGCTTTGTTGAGG | (Lopez-Gomez et al., 2012) |
| *LjNSP2* | F: CATCGACTCCATGATTGACG | (Lopez-Gomez et al., 2012) |
| | R: GGTTGTTGTTGTCGTGGTTG | (Lopez-Gomez et al., 2012) |
| *LjIPN2* | F: GAGCCCTTTGATGTGGAGTGA | |
| | R: GCTCTTCTTGGTGATGGCCT | |
| *LjENOD40-1* | F: GGAGGTATGCTCAAACATTC | (Flemetakis et al., 2000) |
| | R: CCTACTCATCTGCAGAAACTG | (Flemetakis et al., 2000) |

| | | |
|---|---|---|
| *LjENOD2* | F: CAGGAAAAACCACCACCTGT | (Radutoiu et al., 2003) |
| | R: ATGGAGGCGAATACACTGGTG | (Radutoiu et al., 2003) |
| **Quantification of FsK colonization within *Lotus japonicus* root tissues** | | |
| *Fusarium-ITS* | F: TGGTCATTTAGAGGAAGTAA | (Skiada et al., 2019) |
| | R: GGTATGTTCACAGGGTTGATG | (Skiada et al., 2019) |
| *Fusarium-TEF1a* | F: CCCCTCCAGGATGTCTACAA | (Skiada et al., 2019) |
| | R: GGAAGACCCTCAGTGAGCTG | (Skiada et al., 2019) |
| **Lys13/14 RNAi (only the primer sequence is presented; primers were designed with attb sites as described for Gateway cloning, Invitrogen)** | | |
| *Lys13/Lys14* | F: ACCGGCGAGGTTACAGATAG | This study |
| | R: GGCGACGATTGTAGAAACAGA | This study |
| **Verification of the presence of hairpin expressing T-DNA in regenerated hairy roots** | | |
| *pUB-GFP* | F: GCTACCCCGACCACATGAA | |
| | R: GTTGTACTCCAGCTTGTGCC | |
